# *In vivo* lineage conversion of vertebrate muscle into early endoderm-like cells

**DOI:** 10.1101/722967

**Authors:** Clyde Campbell, Joseph J. Lancman, Raquel Espin Palazon, Jonatan Matalonga, Jiaye He, Alyssa Graves, Xin-Xin I. Zeng, Rashmi Mishra, Jan Huisken, David Traver, P. Duc Si Dong

## Abstract

The extent to which differentiated cells, while remaining in their native microenvironment, can be reprogrammed to assume a different identity will reveal fundamental insight into cellular plasticity and impact regenerative medicine. To investigate *in vivo* cell lineage potential, we leveraged the zebrafish as a practical vertebrate platform to determine factors and mechanisms necessary to induce differentiated cells of one germ layer to adopt the lineage of another. We discovered that ectopic co-expression of Sox32 and Oct4 in several non-endoderm lineages, including skeletal muscle, can specifically trigger an early endoderm genetic program in a cell-autonomous manner. Gene expression, live imaging, and functional studies reveal that the endoderm-induced muscle cells lose muscle gene expression and morphology, while specifically gaining endoderm organogenesis markers, such as the pancreatic specification genes, *hhex* and *ptf1a*, via a mechanism resembling normal development. Endoderm induction by a pluripotent defective form of Oct4, endoderm markers appearing prior to loss of muscle cell morphology, a lack of dependence on cell division, and a lack of mesoderm, ectoderm, dedifferentiation, and pluripotency gene activation, together, suggests that reprogramming is endoderm specific and occurs via direct lineage conversion. Our work demonstrates that within a vertebrate animal, stably differentiated cells can be induced to directly adopt the identity of a completely unrelated cell lineage, while remaining in a distinct microenvironment, suggesting that differentiated cells *in vivo* may be more amenable to lineage conversion than previously appreciated. This discovery of possibly unlimited lineage potential of differentiated cells *in vivo* challenges our understanding of cell lineage restriction and may pave the way towards a vast new *in vivo* supply of replacement cells for degenerative diseases such as diabetes.

## Introduction

In animals, from flat worms to humans, nearly all cell lineages develop from one of three germ layers, established during the earliest stages of development. Despite several hundred millions of years of evolution, the type of specialized cells (i.e., neuronal, muscle, or pancreatic cells) that can develop from each of the distinct germ layers (ectoderm, mesoderm, or endoderm, respectively), remain well conserved among highly divergent animals, suggesting that lineage identities are restricted to specific germ layers. This paradigm is consistent with the prevailing interpretation of the Waddington’s epigenetic landscape ^1^ model suggesting that during animal development, a cell’s lineage potential becomes increasingly restricted as they differentiate. Furthermore, lineage restriction within germ layers appears to also apply to induced *in vivo* cell lineage reprogramming of vertebrate cells, as directed lineage transdifferentiation has only been shown among cell types from the same germ layer ^2^. However, in the invertebrate roundworm *Caenorhabditis elegans*, it was demonstrated that two specific non-endoderm cell lineages, can be artificially converted into intestinal cells, which are normally of endoderm origin ^3^. It remains unclear whether *in vivo* intergerm layer lineage conversion is unique to these cell types or to this and other invertebrates ^4^. The ability to directly reprogram a differentiated vertebrate cell identity *in vivo*, without spatial or lineage limitations, would indicate that the developmental origin of a cell and its natural microenvironment does not absolutely limit its potential identity, challenging the dogma of *in vivo* developmental lineage restriction and opening potential new avenues for regenerative medicine.

### *In vivo* platform to identify endoderm inducing factors

To investigate whether stably differentiated vertebrate cells can be lineage converted *in vivo* into unrelated cell types (across germ layers and in distinct microenvironments), we leveraged the zebrafish embryo. This animal model is highly amenable to transgenic modification, and due to its rapid embryonic development, functionally differentiated muscle cells are already present at 24 hpf (hours post fertilization), as indicated by a beating heart and body movements. We aimed to induce endoderm lineages from differentiated, non-endoderm-derived cells and in tissues not closely localized to gut endoderm derived tissues. Unlike differentiated cells originating from the mesoderm and ectoderm which are often intermingled, differentiated cells of the endoderm lineage remain largely contiguous with the gut tube, facilitating the identification of ectopically induced endoderm cells in non-endoderm derived tissues.

In contrast to previous *in vivo* reprogramming strategies to induce a specific endoderm cell type ^5^ or a pluripotent intermediate, our novel strategy is to trigger the early endoderm genetic program. With this approach, we predict that early-endoderm induced cells will have the developmental potential to progress towards various endoderm lineages and ultimately adopt distinct endoderm cell fates. Zygotes were injected with DNA constructs containing candidate endoderm specification genes (individually or in combination) driven by the heat shock inducible *hsp70l* promoter ^6^, allowing for temporally controlled, transient, and spatially mosaic transgene expression throughout the fish (Fig.1A-A”). With this approach, transgene expression levels will vary extensively between cells due to variable transgene dose. Therefore, this *in vivo* platform allows for highly diverse conditions of transgene expression, in a variety of different cell types, and at different states of differentiation, thereby enhancing the chance of discovering factors with lineage conversion potential.

*sox32* and *oct4* were both previously shown to be required during zebrafish gastrulation for normal endoderm lineage specification^7-9^. With induction beginning at 24 hpf, we found that expression of either zebrafish *sox32* (*hsp70l:H2BmCherry-P2A-sox32*) or mouse *Oct4* (*pou5f3*; *hsp70l:mCherryCAAX-P2A-Oct4*) alone failed to significantly induce ectopic expression of *sox17* (*sox17:GFP*), the earliest definitive endoderm marker in fish and mammals ^10^ (Figure 1B-C”). However under the same conditions, co-injection of both constructs resulted in obvious ectopic *sox17:GFP* expression at 48 hpf in multiple regions throughout the embryos, including the myotomes, epidermis, and spinal cord (Figure1D-D”, Supplement Figure 1). Upon closer examination of sox17:GFP-positive cells within the myotomes, we found that most have multiple nuclei and an elongated and striated cell morphology (see Figure 1D’). These cells also coexpressed the transgene reporters for both *sox32* (nuclear mCherry) and *Oct4* (membrane mCherry), indicating that differentiated skeletal muscle cells can be cell-autonomously induced by Sox32 and Oct4 to express *sox17*. Because *sox17* is also expressed in a sub-population of endothelial cells during development, we examined endothelial markers and found no ectopic induction of *fli1a:GFP* or *kdrl:GFP* (Supplement Figure 2A-D’) in muscle cells of the myotome, suggesting that the ectopically induced *sox17* expression does not indicate endothelial identity, but more likely endoderm. Further, forced coexpression of both *sox32* and *Oct4* can also cell-autonomously induce ectopic *foxa3* (*foxa3:GFP*), a gene normally expressed in a broad subset of the endoderm (Figure 1E-E’). Consistent with normal endoderm development, we find that ectopic *foxa3:GFP* expression appeared later than *sox17*. These results indicate that Sox32 and Oct4 can intrinsically induce the earliest genetic markers of the endoderm lineage. Our finding that the combination of mouse (and human; not shown) Oct4 with zebrafish Sox32 can induce *sox17:GFP* and *foxa3:GFP* expression in fish also supports a conserved genetic mechanism for induction of endoderm lineage specification between fish and mammals.

**Figure 1:**
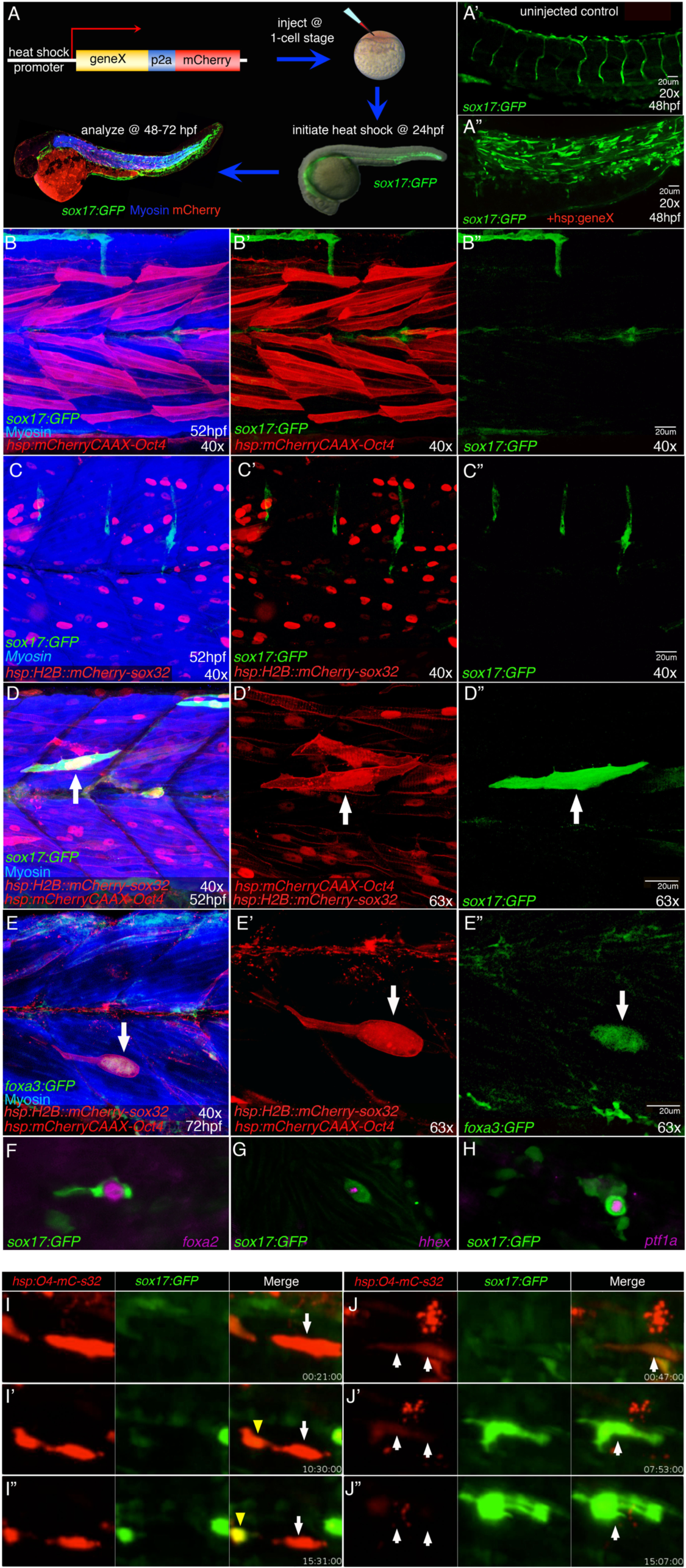
in vivo identification of factors that induce expression of early endoderm factors. **(A)** Experimental platform: transient expression constructs with candidate endoderm inducing genes and *mCherry* fluorescent reporter (nuclear or membrane) under control of the heat shock promoter (hsp70l) were injected into transgenic *sox17:GFP* zebrafish at the 1-cell stage and heat shock induced at 24 hours post fertilization (hpf). Live embryos were screened between 48-72hpf by fluorescent microscopy for ectopic sox17:GFP expression in non-endoderm tissue. **(A’, A”)** Three dimensional (3D) rendering of 48 hpf *sox17:GFP* zebrafish trunk of **(A’)** Uninjected control showing normal *sox17:GFP* expression in the vasculature and the gut endoderm and **(A”)** an injected experimental sample displaying robust induction of sox17:GFP outside of its endogenous domains. **(B-D)** 48 hpf *sox17:GFP* (green) myotomes with labelled Myosin (blue) following heat shock induction of *mCherry-CAAX-P2A-Oct4* **(B, B’ membrane-Red)** or *H2B::mCherry-P2A-sox32* **(C, C’ nuclear-Red)** individually or together **(D, D’, E, E’ membrane and nuclear-Red)**. Neither *Oct4* **(B-B”)** nor *sox32* **(C-C”)** alone significantly induced ectopic *sox17:GFP* expression (green). **(D-D”)** Co-expression of both *sox32* and O*ct4* **(membrane and nuclear red; arrow)** can lead to cell autonomous induction of ectopic sox17:GFP expression in myocytes **(arrow, D”). (E-E”)**, Co-expression of O*ct4* and *sox32* can also cell autonomously activate the expression of another endoderm marker, *foxa3:GFP (green)*, in 72 hpf myocytes **(arrow, E”). (F-H)** Whole mount *in situ* hybridization of s*ox17:GFP* embryos injected with *hsp:Oct4-P2A-mCherry-P2A-sox32* to assess co-expression with foregut endoderm genes. Putative muscle cell showing co-expression of sox17:GFP protein (green) with mRNA expression (magenta) of *foxa2* **(F)**, *hhex* **(G)**, and *ptf1a* **(H). (I-J”)**. Selected live image movie stills of individual myocytes with transgene expression (Red; *hsp:Oct4-P2A-mCherry-P2A-sox32)* in transgenic *sox17:GFP* (Green) zebrafish from 48-72hpf using light sheet confocal microscopy. **(I-I”)** Arrow(s) point to an individual myocyte that splits into two presumptive cells in about 15 hours with the cell on the left (yellow arrow) exhibiting an increase in *sox17:GFP* expression and adopting a stellate shape (Supplement Movie #2). **(J-J”)** Arrows point to a single cell rapidly changing color (red to green) and shape in less than 15 hours (Supplement Movie #3).

**Figure 2:**
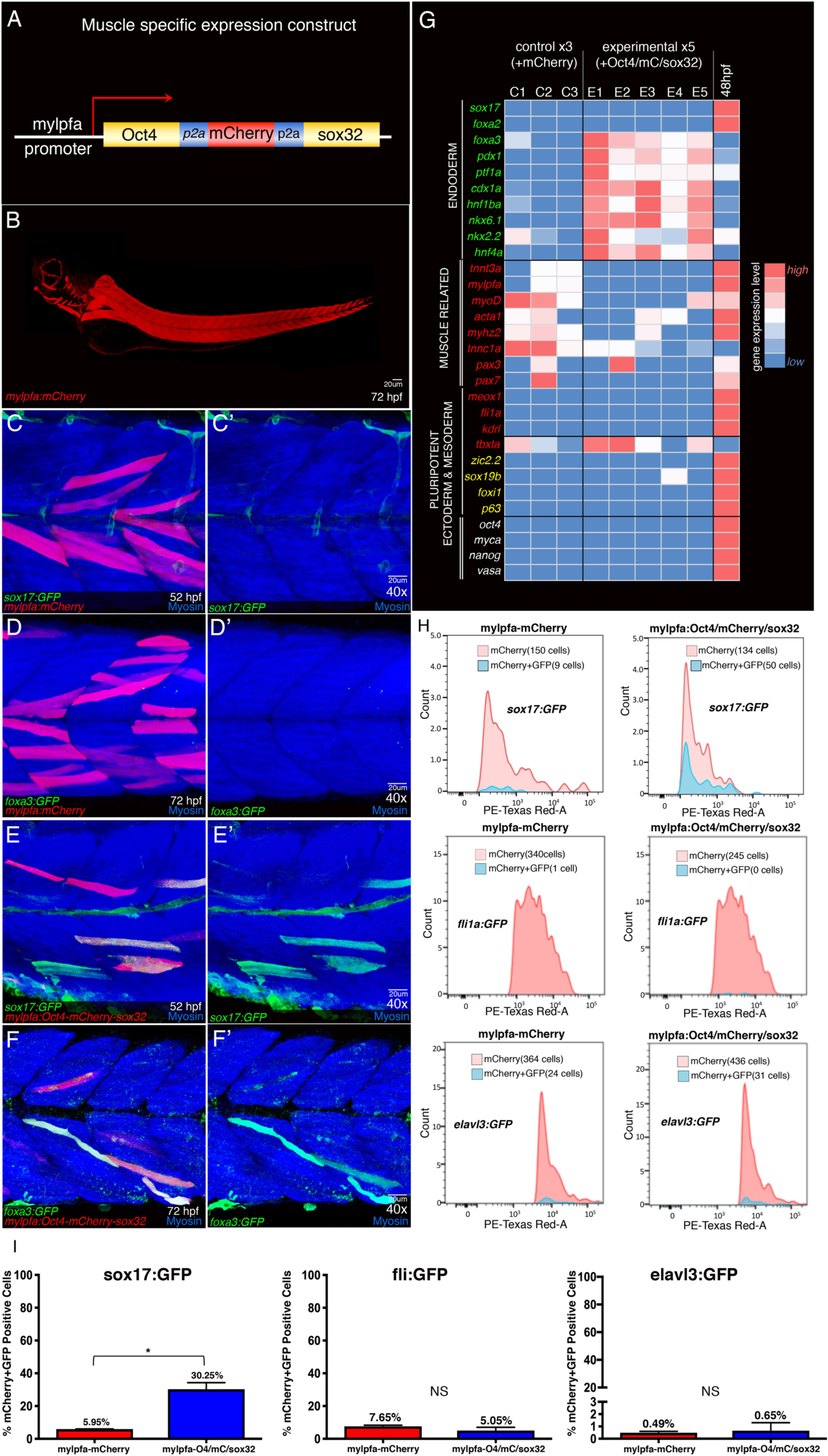
Oct4 and sox32 specifically triggers the endoderm genetic program, while inhibiting muscle identity. **(A)** Diagram of DNA construct to restrict expression of factors to fast muscle cells using the tissue specific promoter *mylpfa* to drive expression of the polycistronic transgenes (*Oct4-P2A-mCherry-P2A-sox32*). **(B)** Embryo at 72 hpf injected with the same *mylpfa* promoter driving only *mCherry* (*mylpfa:mCherry*) appears to show exclusive expression **(red)** in differentiated fast muscle cells despite broad transgene expression (n=5/5). **(C-F’)** Confocal z-stacks of 52 hpf *sox17:GFP* and 72 hpf *foxa3:GFP* myotomes labelled with Myosin **(blue)**. Injected with *mylpfa:mCherry*, controls show no GFP induction in muscle cells expressing mCherry **(red-C, C’; D, D’)**. In contrast, with *mylpfa:mOct4-P2A-mCherry-P2A-sox32* injected, sox17:GFP **(E, E’; green)** and foxa3:gfp **(F, F’; green)** are expressed in mCherry positive myocytes **(Blue, E-F’). (G)** Heat map summarizing qPCR analyses of lineage markers in 48hpf FAC sorted mCherry positive myocytes from *mylpfa:mCherry* (3 biological replicates) and *mylpfa:oct4-P2A-mCherry-P2A-sox32* (5 biological replicates) injected embryos and 48hpf whole embryos (technical control). **Red**, Higher expression; **Blue**, Lower expression. Endoderm lineage and organogenesis (green text) genes are upregulated, whereas muscle (red text) genes are downregulated. Mesoderm (red text), ectoderm (yellow text), and pluripotency (white text) genes are not upregulated. **(H)** Representative histograms of FAC analyzed mCherry positive and mCherry/GFP double positive myocytes from embryos injected with either *mylpfa:mCherry* (left column) or *mylpfa:Oct4-P2A-mCherry-P2A-sox32* (right column) to assess induction efficiency of *sox17:GFP* (TOP), *Fli1:GFP* (MIDDLE), and *elavl3:GFP* (BOTTOM). Red peaks represent all cells and blue peaks represent only mCherry/GFP double positive cells. **(I)** Quantification of the number of mCherry/GFP double positive cells from histogram data shown in H: **(Left)** mCherry**/**sox17:GFP, **(Middle)** mCherry/fli:GFP and **(Right)** mChery/*elavl3:GFP* embryos. Shown are the means+/-SEM from two separate, independent experiments. *P=0.026 for sox17:GFP cells by unpaired, two-tailed t-test. fli:GFP and elavl3:GFP differences were NS.

To increase the number of cells expressing both Oct4 and Sox32, the coding sequence for each factor, in addition to mCherry, were placed within the same polycistronic construct, downstream of the *hsp70l* promoter (*hsp70:Oct4-P2A-mCherry-P2A-sox32*). Using this expression construct, we found ectopic *sox17:GFP* in p63 expressing cells (epidermal) and Elavl3/4 expressing cells (neuronal) suggesting that cells originating from the surface ectoderm and neural ectoderm, respectively, can be induced by Sox32 and Oct4 to express *sox17* (Supplement Figure 1A-B”). To further investigate *in vivo* genetic and cellular changes induced by ectopic expression of these genes, we focused our analysis on skeletal muscle cells in the trunk of the animals. Differentiated skeletal muscle cells can be identified based on their elongated shape, striations, and multiple, regularly spaced nuclei, and are abundantly found in a consistent, parallel pattern spanning individual myotomes. The appearance of ectopic *sox17*:GFP and *foxa3*:GFP expression in muscle cells with these characteristics suggests that differentiated cells are indeed being induced to activate expression of these early endoderm genes, and further suggests that a complete loss of muscle morphology and cell division is not a prerequisite for their induction. However a subset of skeletal muscle cells expressing *Oct4* and *sox32* transgenes do exhibit variable loss of muscle morphology, including partial or complete loss of striation and elongated cell shape, and adopting a more stellate shape (Figure 1E and J). Interestingly, the nuclei in reporgrammed muscle cells appear to aggregate together towards the center of the cell (n=20/22 cells; Supplement Figure 3), reminiscent of striated muscle regeneration ^11^. Live imaging from 48 to 72 hpf, using light sheet microscopy, shows highly dynamic changes in cell morphology (Supplement Movie 1). In particular, some muscle cells appear to be separating into two individual cells, potentially becoming mononucleated (Figure 1I-I”; white and yellow arrows, Supplement Movie 2). Further, we observed that cells with induced *sox17*:GFP form cellular extensions in multiple directions. Intriguingly, mCherry expression can be rapidly cleared, suggesting an active process for protein turnover in cells that are being reprogrammed (Figure 1J-J”; double arrows, Supplement Movie 3). Consistent with the rapid clearance of transgenic mCherry protein lost, endogenous Myosin proteins (and transcripts), which are normally abundantly expressed, can become undetectable in certain reprogrammed muscle cells (Supplement Figure 4). Rapid turnover of structural proteins such as Myosin would explain the rapid loss of the skeletal muscle morphology, as these cells are dense with structural proteins necessary for their elongated shape and striations.

**Figure 3:**
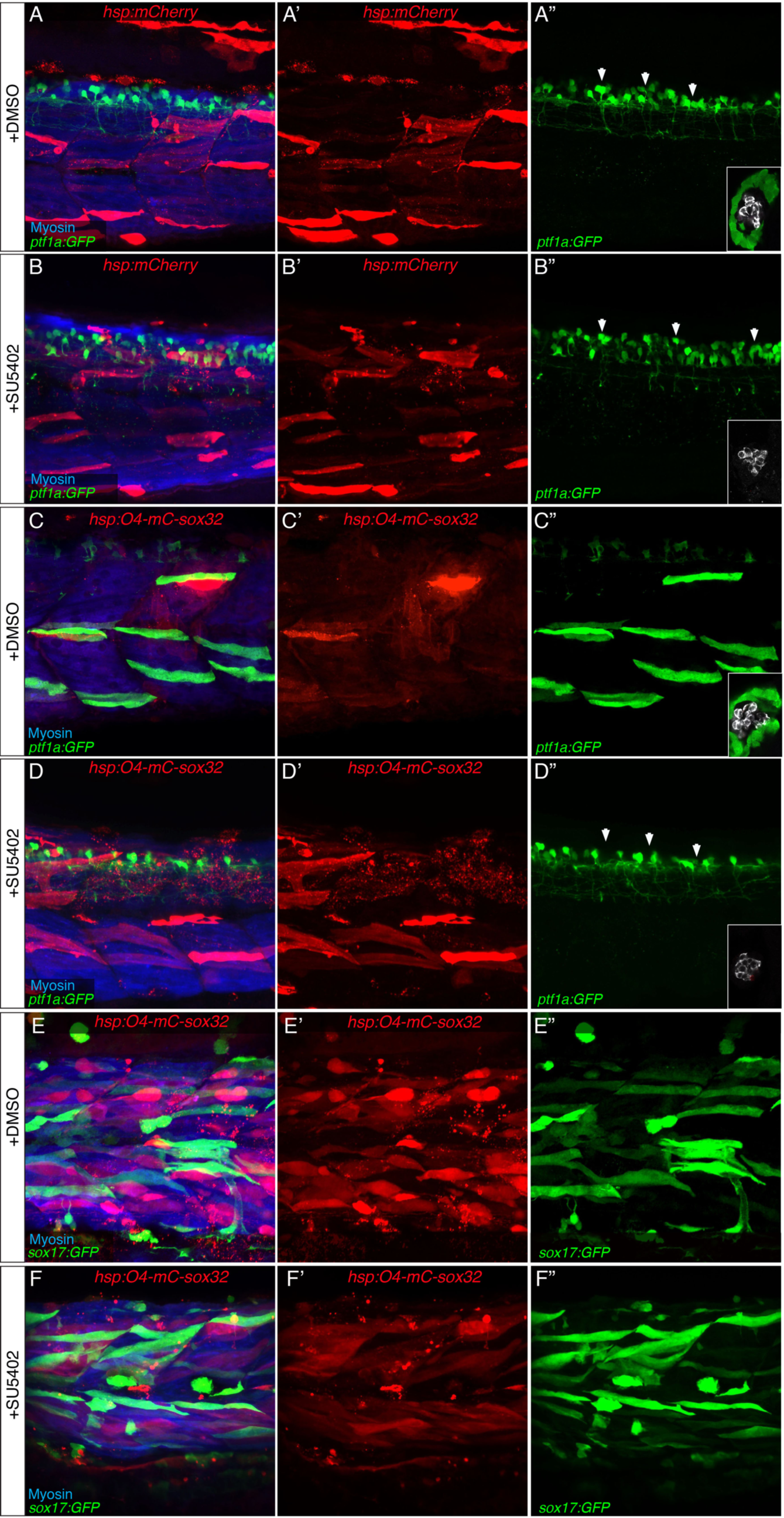
ptf1a expression in endoderm-induced muscle cells requires FGF signaling. 3D rendering of 72 hpf *ptf1a:GFP* zebrafish myotomes from embryos injected with either *hsp:mCherry* or *hsp:Oct4-P2A-mCherry-P2A-sox32* and treated with either the vehicle DMSO **(A, C)** or the FGFR kinase inhibitor SU5402 **(B, D)**. DMSO treatment did not affect reprogramming outcomes in the *ptf1a:GFP* background following induction of control *hsp:mCherry* **(A-A”: +ectopic ptf1a:GFP; n=0/118)** compared to SU5402 treated controls **(B-B”: n=0/106)**. Note that DMSO also has no effect on normal exocrine pancreas development, marked by *ptf1a:GFP* **(green; inset A”, C”)** or pancreatic endocrine beta cell development, labelled with Insulin **(white; inset A”, C”)**. DMSO treatment did not impact reprogramming efficiency with hsp:*Oct4-P2A-mCherry-P2A-sox32*, in ptf1a:GFP transgenics **(C-C”: n=27/121).** However, SU5402 did strongly inhibit *ptf1a:GFP* induction in myocytes expressing Oct4 and Sox32 **(D-D”:n=0/113)**. Similarly, su5402 also inhibits ptf1a-GFP induction during normal exocrine pancreas development **(inset B”, D”)** but it does not affect endocrine pancreas beta cell development as marked by Insulin **(white; inset B”, D”)** or ptf1a:GFP expression in neural tissue **(arrow heads A”, B”)**. Importantly, SU5402 does not affect *sox17:GFP* induction by Oct4 and sox32 in reprogramed myocytes **(green; +ectopic sox17:GFP n=51/88)** as sox17:GFP induction is similar in DMSO treated animals (**green; E-E”: n=56/94)**.This is consistent with normal development in which Fgf signaling is not required for early induction of sox17 expression but is necessary for pancreatic ptf1a expression.

**Figure 4.**
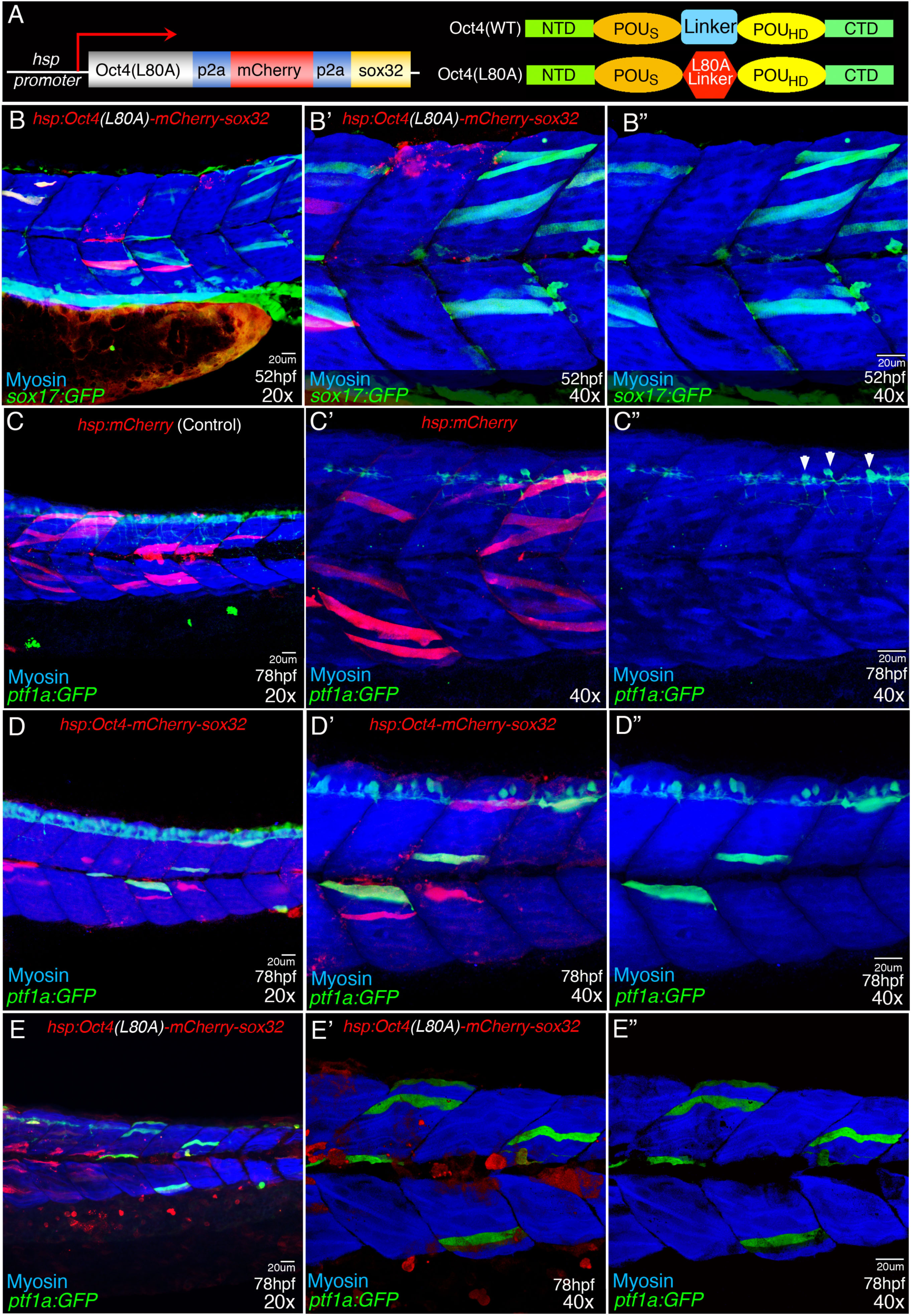
Pluripotency defective Oct4 can induce endoderm genes in muscles. **(A)** Schematic of expression construct containing the pluripotent defective mutant Oct4 (left) and a diagram comparing the structure of wild type Oct4 protein **(right, Top)** to the pluripotent defective L80A Oct4 variant with a missense mutation in the linker region between POU domains **(right, Bottom). (B-E”)** 3D rendering of 52hpf *sox17:GFP* 78hpf and *ptf1a:GFP* zebrafish myotomes labeled with Mysoin (Blue) to mark myocytes in zebrafish injected with either *hsp70:Oct4(L80A)-P2A-mCherry-P2A-sox32* which contains the pluripotent defective mutant Oct4 **(B-B”, E-E”)**, *hsp:mCherry* **(C-C”) or** h*sp70:Oct4-P2A-mCherry-P2A-sox32* **(D-D”)** or Ectopic ptf1a:GFP (green) expression is never detected outside of the dorsal spinal cord neurons following overexpression of mCherry alone **(B-B”; Arrowheads: n=0/30)**. However, expression of sox32 and either wild type Oct4 **(C-C”)** or the L80A Oct4 mutant **(D-D”: n=12/52) can** both induce *ptf1a:GFP* (green) in myocytes **(myosin-Blue)**. Co-expression of sox32 and the L80A Oct4mutant can also induce *sox17:GFP* in myocytes at 52hpf **(green, E-E”: n=11/30)**.

To investigate whether induced muscle cells with *sox17*:GFP expression function like early endoderm and progress in a development-like manner, we examined later markers of endoderm regionalization and organogenesis. Endogenous endoderm *sox17* mRNA expression is temporally restricted to a short time window before the end of gastrulation (∼7-10 hpf) and is therefore not coexpressed with later endoderm genes ^12^. Consistent with the early and brief duration of *sox17* transcript expression observed in normal endoderm development, ectopic *sox17* mRNA expression was limited to within the initial several hours following heat shock induction of the *Oct4* and *sox32* transgenes (not shown). However, perdurance of GFP allows for examination of *sox17*:GFP-positive cells beyond the loss of ectopic *sox17* transcript expression. By 48 hpf, *sox17*:GFP-positive cells in the myotome can be found to coexpress transcripts of the foregut endoderm gene *foxa2* and pancreas specification genes, *hhex* and *ptf1a* (Figure 1F-H). In contrast, transcripts specific to muscles such as *myh7* (myosin heavy chain 7) mRNA are lost from most *sox17*:GFP positive muscle cells (Supplement Figure 4A-B”). These findings suggest that ectopic Oct4 and sox32 expression can inhibit muscle specific transcripts while triggering the endoderm developmental genetic program.

### The early endoderm genetic program is specifically triggered while muscle gene expression is lost

To express transgenes in a specific type of differentiating skeletal muscle cell, the *mylpfa* (*myosin light chain, phosphorylatable, fast skeletal muscle a*) promoter ^13^ was used to drive Oct4 and Sox32 expression (Figure 2A). With a construct containing the *mylpfa* promoter driving mCherry alone, we confirmed that the activity of this promoter is almost exclusively restricted to fast muscle cells (and not found in non-skeletal muscle cells) in injected embryos (Figure 2B) and does not induce expression of early endoderm flourescent reporters (Figure 2C-D’). With *Oct4, sox32*, and *mCherry* all driven by this lineage specific promoter (*mylpfa:Oct4-P2A-mCherry-P2A-sox32*), we found ectopic *sox17:*GFP and *foxa3:*GFP induced only in the fast skeletal muscle layer of the myotomes (Figure.2E-F’), but not in other tissues. Flow cytometry analysis of mCherry positive cells from these injected embryos at 48hpf reveal that approximately 30% of muscle cells expressing *sox32/Oct4* also ectopically express *sox17*:GFP (Fig.2H-Top, 2I-Left). Given that *sox17*:GFP positive cells continue to arise after 48hpf and that mCherry expression can also be rapidly lost (see Figure1J-J”, Supplement Movie 1), the actual level of efficiency is likely higher – albeit efficiency does vary significantly among individual embryos. In contrast, under the same experimental conditions, no significant induction of endothelial marker *fli1a*:GFP or neural marker *elavl3*:GFP was detected, consistent with specific induction of early endoderm transcripts (Figure 2H-Middle; Bottom, 2I Middle, Right, Supplement Figure 2C-F’).

To more broadly assess transcriptional changes induced in reprogrammed muscle cells, we pooled sorted mCherry-positive cells from fish injected with either the *mylpfa* promoter driving *mCherry* alone (control) or together with *Oct4* and *sox32*, and carried out qPCR analysis. Consistent with our histological studies using fluorescent reporter lines and whole mount *in situ* hybridization, early and late endoderm genes, *sox17, foxa2, hnf1ba*, hnf4a, and *ptf1a* mRNA expression are all significantly upregulated in sorted muscle cells with *sox32* and *Oct4* transgene expression (Figure 2G, Supplement Figure 5). These results suggest that in fast muscle cells, the endoderm specification genes *sox32* and *Oct4* can trigger genetic programs for multiple endoderm lineages. We also examined markers of other germ layers to determine the specificity of the endoderm induction. Mesoderm genes, *tbxta* (*T*; *brachyury*) and *meox1*, surface ectoderm genes, *foxi1* and *p63*, and neural ectoderm genes, *sox1a, zic2.2*, and others, are not upregulated, suggesting neither mesoderm or ectoderm lineages are induced (Figure 2G). However, we found that the muscle genes *myod, myhz2, tnnt3a*, and *mylpfa* are downregulated, consistent with the loss of muscle cell morphology and myosin protein/mRNA expression described above. Together, these gene expression studies indicate that forced expression of sox32 and Oct4 in fast skeletal muscle cells leads to a loss of the muscle genetic program and to a specific activation of the endoderm genetic program.

### Induced endoderm cells proceed through a developmental mechanism to express pancreatic gene *ptf1a*

To explore the mechanism by which the induced endoderm (iEndo) cells proceed to express later organogenesis genes, we focused on reprogrammed muscle cells with ectopic expression of the pancreas specification gene *ptf1a*. Misexpression of *sox32* and *Oct4* can lead to muscle cells with *ptf1a*:GFP by 48-72 hpf in about one in five injected embryos. iEndo muscle cells with ectopic *ptf1a*:GFP are most often found in the posterior third of the fish. The low frequency and spatial propensity of induced *ptf1a*:GFP expression led us to posit that only certain iEndo cells with optimal intrinsic (intracellular) and extrinsic (extracellular/microenvironment) conditions will proceed toward a specific endoderm lineage pathway such as a liver, intestine, or pancreatic genetic program. The skeletal muscle microenvironment has previously been demonstrated to be permissive for human pancreas cell differentiation and function ^14-17^. Because neuronal genes are not induced, the ectopic *ptf1a* expression observed is unlikely to represent the normal cerebellar or retinal domains of *ptf1a* expression. Furthermore, because iEndo muscle cells with ectopic *ptf1a* expression can be found to coexpress *sox17*:GFP (see Figure 1H), they are more likely to be pancreatic endoderm. However, a lack of late pancreatic genes in these cells indicates that conditions are suboptimal for further differentiation.

The temporal sequence of endoderm genes expressed in iEndo cells appears to recapitulate that of endogenous endoderm development, leading us to functionally assess whether the ectopic *ptf1a* expression occurred via a genetic pathway analogous to normal development. FGF signaling was shown to be necessary for foregut endoderm cells to adopt ventral pancreatic endoderm lineage in normal zebrafish development and in human stem cells differentiation models ^18,19^. Further, FGF signaling is highest posteriorly where ectopic *ptf1a*:GFP-positive muscle cells are most often observed ^20^. Blocking FGF receptor tyrosine kinase signaling with the inhibitor SU5402 prevents iEndo muscle cells and endogenous foregut endoderm from expressing *ptf1a*:GFP but does not prevent ectopic *sox17:*GFP (Figure 3A-F”). These findings suggest that iEndo cells do not spontaneously express *ptf1a*, but rather proceed through a step-wise genetic program resembling normal, Fgf signaling dependent, pancreas development.

### Pluripotency is not required for *in vivo* induced endoderm

Examples of *in vivo* cell lineage plasticity, including natural transdifferentiation in worms, fin regeneration in zebrafish, and maintenance of the neural crest lineage potential in frogs, have implicated a pluripotency mechanism ^21-23^. These findings led us to ask whether our induction of muscle into endoderm lineage conversion using Oct4 and Sox32 also involves a pluripotency mechanism. The lack of induction of the skeletal muscle progenitor genes *pax3* and *pax7* (Figure 2G), suggests that lineage conversion does not involve the dedifferentiation of skeletal muscle cells. The appearance of endoderm gene expression prior to loss of muscle morphology suggests that cell division is not required for iEndo cells. Consistently, inhibition of cell proliferation with aphidicolin does not prevent the induction endoderm transcripts by *sox32* and Oct4 (not shown). Together with a lack of mesoderm and ectoderm gene activation, we posit that lineage conversion of muscle to endoderm does not involve a pluripotent intermediate. Consistently, qPCR expression analysis does not show upregulation of standard pluripotent mRNA markers, including endogenous *oct4, myca, vasa*, or *nanog* (Figure 2G). However, it may be possible that Oct4 functions as a pluripotent factor without requiring the iEndo cells to have gone through a detectable pluripotent intermediate. Mammalian Oct4, in combination with other Yamanaka factors, was previously shown to be able to reprogram fish cells in culture to pluripotency ^24^. To assess the requirement of Oct4’s pluripotency function for reprogramming muscle into endoderm *in vivo*, we used a modified form of Oct4 which has an amino acid substitution in the linker domain, previously shown to be transcriptionally active but unable to induce pluripotency (Oct4(L80A); Figure 4A) ^25^. As with wild-type Oct4, mis-expression of mutant Oct4(L80A) with Sox32 (*hsp70:Oct4(L80A)-P2A-mCherry-P2A-sox32*) can induce muscle cells to express *sox17*:GFP, as well as lead to cell shape changes (Figure 4B-B”). Moreover, like wild type Oct4 iEndo cells, these *Oct4(L80A)* iEndo cells can also proceed to exhibit *ptf1a*:GFP expression and lose myosin (Figure 4C-E”, Supplemental Figure 3F-F”), suggesting that they can progress towards a pancreas developmental genetic program, while rapidly losing muscle protein expression. These findings demonstrate that a robust pluripotency mechanism via Oct4 is not required to induce muscle cells into early endoderm-like cells, further supporting our conclusion that *in vivo* lineage conversion across germ layers can be induced directly, independent of a pluripotent intermediate. This finding also functionally demonstrates for the first time, that Oct4’s roles in endoderm specification may be distinct from its well defined role in pluripotency.

## Discussion

Within a vertebrate embryo, we demonstrate that differentiated mesoderm-derived skeletal muscle cells can be induced with ectopic expression of just two transcription factors, Sox32 and Oct4, to cell-autonomously trigger early endoderm-like development. These early endoderm-induced cells can rapidly lose muscle cell morphology and gene expression while transitioning through an endoderm genetic program resembling normal endoderm development. This lineage conversion process appears to be direct as no evidence was found to support a pluripotent or dedifferentiated intermediate state. As with normal endogenous pancreatic *ptf1a* expression, expression of *ptf1a* in iEndo muscle cells also requires Fgf signaling. Expression of *ptf1a*, in addition to other endoderm organogenesis specification factors, shows that these iEndo cells can function to give rise to diverse endoderm lineage genetic programs. Expression of organogenesis markers also suggests the potential for these iEndo to be further coaxed, with additional factors, into a specific functional differentiated endoderm organ cell type such as pancreatic beta-cells, which would be useful for diabetics.

Waddington’s epigenetic landscape model suggests that as cells differentiate during normal development, they become more restricted from adopting other lineage identities. However, requiring only a few transcription (intrinsic) factors, transdifferentiation across germ layers using *in vitro* approaches has proven to be surprisingly easier than the Waddington model would predict, challenging this paradigm of limited lineage potential ^26^. Yet, with *in vitro* lineage reprogramming, removal of the cells from their native microenvironment and exposing them to artificial culture conditions may compromise their lineage stability, facilitating their reprogramming. Importantly, Waddington’s model addresses cell lineage constraint in the context of an embryo, where a cell’s normal microenvironment may also be restricting its lineage potential. Our work, using zebrafish embryos, demonstrates that despite the differentiated muscle cells remaining in their native microenvironment, ectopic expression of only two transcription factors is able to repress muscle identity while triggering the early endoderm genetic program, suggesting that both intrinsic and extrinsic factors maintaining muscle cell identity can be surmounted to induce conversion towards an unrelated lineage identity. Because we observe that ectopic *sox17* expression is also induced by Oct4 and Sox32 in other differentiated, non-muscle, cell types in other regions of the embryo, it is likely that other cell lineages, in other distinct microenvironments, are also amenable to induced *in vivo* lineage conversion.

Analogous to the commonly used MEFs (mouse embryonic fibroblasts) for *in vitro* lineage reprogramming studies, differentiated cells within the zebrafish embryo were used here as a practical vertebrate *in vivo* discovery platform to identify lineage reprogramming factors and investigate their mechanisms. This *in vivo* platform allows for transient and mosaic expression of transgenes, at greatly varying levels, in a large number of cells within a large number of animals, thereby increasing the likelihood of discovering combinations of factors capable of inducing lineage conversion. Testing reprogramming factors on differentiated cells throughout the embryonic fish has key advantages. The rapid development and transparency of the zebrafish embryo together with transgenic fluorescent lineage reporter lines allows for quick assessment of candidate reprogramming factors in live animals. In contrast to MEFs, the particular cell lineage reprogrammed, and its specific microenvironment, can be more definitively identified, allowing for the assessment of how intrinsic and extrinsic factors influence the lineage conversion process. But similar to MEFs, differentiated cells at developmental stages are presumably more amenable to cell lineage reprogramming, providing a ‘sensitized setting’ for revealing cell lineage reprogramming factors that would otherwise be difficult to uncover. Additional factors, including epigenetic modifying small molecules, may be necessary for reprogramming more mature or aged differentiated cells, as previously shown in adult mice ^27^.

The high variability of reprogramming efficiency among individual embryos indicates great potential in the number of differentiated cells that are amenable to induction by Oct4 and sox32 to initiate the endoderm program. Conversely, the high variability of reprogramming efficiency also indicates that there are factors and conditions yet be uncovered that may allow for more consistent and efficient induction of lineage conversion. This efficiency hurdle will need to be overcome to yield enough cells for single-cell systems analyses. Uncovering these factors to enhance lineage reprogramming will undoubtedly lead to further mechanistic insight into direct lineage conversion, both *in vivo* and *in vitro*. Testing additional reprogramming factors may also allow for induction of a distinct endoderm lineage such as pancreatic beta-cells, which will potentially have biomedical applications. Transplantation of replacement beta-cells into various locations in the body ^28^, including in skeletal muscles in humans ^16,17^ suggests that beta-cells can function outside their normal microenvironment to help maintain blood glucose levels. The ability to induce cells outside the foregut to adopt endoderm-like identity, as demonstrated here, is a significant step towards ultimately generating replacement beta-cells from potentially any cell type directly in the body of diabetics. Direct *in vivo* lineage conversion to generate replacement cells ^29,30^ may bypass safety ^31^ and efficacy risks associated with transplantation of *in vitro* engineered pluripotent cells ^32-36^. An unrestricted ability for directly reprogramming any differentiated vertebrate cells *in vivo* into any cell type, in any microenvironment, would greatly expand the potential therapeutic applications of direct, induced *in vivo* lineage conversion for regeneration medicine.

## Materials and Methods

### Animal husbandry

Adult zebrafish and embryos were cared for and maintained under standard conditions. All research activity involving zebrafish was reviewed and approved by SBP Medical Discovery Institute Institutional Animal Care and use Committee (IACUC: UO1 DK105554; Expires 01/18/2021) in accordance with Public Health Policy regarding care and use of laboratory animals and comply with the Guide for the Use of laboratory Animals and the regulations set forth in the Animal Welfare Act and other applicable federal, state and local laws, regulations and policies. The following transgenic lines were used:*Tg(sox17:GFP)*^*s870* 37^, *Tg(gut:GFP)*^*s854* 38^, *Tg(ptf1a:GFP)*^*jh1* 39^, *Tg(elavl3:GFP* ^*knu3* 40^; *Tg(kdrl:EGFP)*^*s843* 41^, *Tg(fli1a:EGFP)*^*y1* 42^.

### Transient Injections and Heat shock

For all experiments, 0.5-1.0nl of plasmid was injected at the 1-cell stage to deliver the following amounts of plasmid DNA:

*hsp:H2B::mCherry-P2A-sox32* and *hsp:mCherryCAAX-P2A-Oct4* (20ng)

*hsp:Oct4-P2A-mCherry-sox32:* (30-35ng)

*hsp:mCherry*: (30ng)

mylpfa:Oct4-P2A-mCherry-sox32:(30-35ng)

*mylpfa:mCherry: (30ng)*

### Antibody Staining

Antibody staining was performed as previously described ^43^ using the following antibodies and staining reagents: (see Supplement Materials and Methods). Samples were imaged on a Zeiss LSM710 running Zen 2010 (Black). Final image processing was performed using Image-J (vs. 1.48b) and/or Photoshop CS3.

### Whole mount In situ hybridization

*In situ* hybridization was performed as previously described^44^, D. LZIC regulates neuronal survival during zebrafish development. Dev. Biol. 283, 322–334, 2005). Samples were imaged on a Leica M165FC using Leica Application Suite v. 3.8 or on a Zeiss Confocal LSM710 running Zen 2010 (Black). Adobe Photoshop CS3 and/or ImageJ64 (vs.1.48b) was used for final image processing. In situ probe plasmid references or primers?

### Fluorescent Whole mount In situ hybridization

*Fluorescent in situ hybridization* combined with immunofluorescence was performed according to published protocols^45-47^ with minor modifications. Probe against *vmhc* was previously described (ZDB-GENE-99123-5) ^48,49^. Following hybridization, embryos were incubated with both anti-digoxigenin-POD (Roche) and chick anti-GFP antibody. *In situ* probe was first detected with TSA Plus fluorescein (Perkin Elmer) followed by incubation with AlexaFluor 594 donkey anti-chick secondary antibody (Invitrogen) to detect *sox17:GFP* localization. Stained embryos were mounted in SlowFade Gold anti-fade reagent with DAPI (Molecular Probes) prior to imaging on a Zeiss Confocal LSM710 running Zen 2010 (Black).

### Heat Shock

5 cycles of heat shock were performed from 24-48hpf. (One cycle=3 hours +heat shock at 38.5°C, 2.5 hours at 28°C)

### Live Imaging

#### Microscope setup

Zebrafish embryos were imaged with a custom-built light sheet microscope (Huisken et al, 2007; PMID#17767321). A laser engine (Toptica MLE, 488nm,561nm) was used as the excitation light source. The fluorescence signal was captured by a water dipping objective (OLYMPUS, UMPLFLN 20XW, 20X/0.5) placed perpendicular to the light sheet direction. Collected signal was filtered (Chroma ET BP525/50, ET LP575) prior to image acquisition.

#### Sample embedding

Zebrafish embryos were de-chorionated at 48hpf and transferred into a low-melting agarose (0.6%) solution prepared with E3 medium and tricaine (Maintained at 37°C). Using a syringe and needle, samples were drawn into a cleaned FEP tube (inner diameter: 0.8mm; wall thickness: 0.4mm; Bola) and the bottom of the tube plugged with solidified 2% agarose for additional support during imaging. The plugged FEP tube was mounted on the stage assembly so that the sample was positioned vertically at the intersection of illumination and detection optical paths of the light sheet microscope. The sample chamber temperature was maintained at 28°C using a custom-built perfusion based temperature control system.

#### Time-lapse acquisition

Samples were moved along the detection axis while being illuminated by the excitation light sheet. Images were taken every 2 microns with a high speed sCMOS camera (Zyla 4.2 PLUS, Andor) at 100 frames per second. The sample was imaged every minute and different excitation wavelengths were imaged sequentially. A typical z-stack of around 300 images was required to cover the region of interest. The overall length of the time lapse recording was approximately 16 hours, resulting in 960 individual 3D stacks in each channel.

### Fluorescent Activated Cell Sorting

To isolate single cells, fluorescence activated cell sorting (FACS) was performed on wild type embryos injected at the 1-cell stage with either *mylpfa:mCherry* or *mylpfa:Oct4-P2A-mCherry-P2A-sox32*. At 24hpf, injected embryos were placed in 0.003% phenylthiourea (PTU) to inhibit melanocyte formation and prevent pigmentation. At 48hpf, embryos were inspected and only healthy/developmentally normal embryos were manually dechorionated. Following collection in 1.5ml eppi tubes, pooled embryos were washed in 1xPBS, incubated in 1xPBS(+Mg/Ca;)+ 0.05 mg/ml Liberase TM (Roche) at 37°C for 60 min and triturated with a P1000 pipette. The resulting suspension was filtered with a 30μm Celltrics cell strainer (Sysmex), spun down (300g for 10min at 4°C) and resuspended in ice cold 1xPBS +0.9% FBS(Gibco). SYTOX Red (Thermo Fischer Scientific) was added at 1:1000 to exclude dead cells immediately prior to sorting for either qPCR analysis or quantification of induction efficiency.

### Quantitative PCR

For qPCR experiments, cells were sorted with a FACSAria II (BD Biosciences). Following FACS, sorted cells were homogenized with a QIAshredder (QIAGEN)and total RNA extracted using the RNeasy Mini Kit (Qiagen). cDNA was synthesized using i-Script Supermix (Bio Rad) and qPCR performed using iQ SYBR Green Supermix (Bio Rad) according to manufacturer protocols. Samples were loaded on a 384 well plate and analyzed on an ABI 7900HT (Applied Biosystems). Primers to detect zebrafish transcripts were designed and are described in Supplement Material and Methods. Relative expression levels of genes were calculated by the following formula: relative expression = 2−(Ct[gene of interest]−Ct[housekeeping gene]). To test statistical significance, the non-parametric Mann-Whitney test was performed (2-tailed, 95% confidence interval).

### Reporter Induction Efficiency

To analyze reporter induction efficiency, *sox17:GFP, elav3:GFP*, or *fli1a:GFP* embryos were injected with either *mylpfa:mCherry* or *mylpfa:Oct4-P2A-mCherry-P2A-sox32*. Upregualtion of fluorescent reporters was quantified by performing FACS using a BD LSRFortessa X-20 (BD Biosciences). Data was analyzed with FlowJo V-10 (FlowJo, LLC). Percent induction efficiency was determined using the following formula: (total number of GFP+ and mCherry+ Cells)/(total number of mCherry+ cells) x 100. Error was determined using the percentage of GFP+ and mCherry+ cells obtained from each respective reporter line injected with *mylpfa:mCherry*.

### SU5402 treatment

3μM SU5402 (sc-204308; Santa Cruz Biotechnology) was added to egg water with 1%DMSO and incubated 24-48 hpf.

### Expression Constructs

Cloning details and sequence available upon request for the following: *Hsp:Oct4-P2A-mCherryCAAX;Alpha-Crystallin:dsRed, Hsp70:H2B::mCherry-P2A-Cas32; Cmlc2:dsRed, Hsp70:Oct4-P2A-mCherry-P2A-sox32; Alpha-Crystallin:dsRed, Hsp70:Oct4(L80A)-P2A-mCherry-P2A-sox32; Alpha-Crystallin:dsRed, Hsp70:mCherry; Alpha-Crystallin:dsRed, mylpfa:Oct4-P2A-mCherry-P2A-sox32; Alpha-Crystallin, mylpfa:mCherry; Alpha-Crystallin.* Oct4 was cloned from FUW-OSKM, a gift from Rudolf Jaenisch (Addgene plasmid # 20328; http://n2t.net/addgene:20328;RRID:Addgene_20328) ^50^.

### Statistics

Statistics and graphs were generated using Prism software (vs.8) Bioinformatics analysis was performed with help from Dr. Jun Yin in the Sanford Burnham Prebys Bioinformatics Core.

## Supporting information

Combined Supplement Figures and Methods

## Author Contributions

P.D.S.D. conceptualized project, designed initial experiments, and oversaw all studies.

P.D.S.D., J.J.L., and C.C. designed experiments. J.J.L. generated all constructs.

C.C., J.J.L., R.E.P., J.M., XX.I.Z., and R.M. performed experiments and analyzed the data with P.D.S.D. A.G. prepared live samples for light-sheet imaging.

J. He built the light-sheet microscope and performed time-lapse image acquisition and processing. J. Huisken oversaw live light-sheet imaging experiments.

D.T. oversaw studies by R.E.P. J.J.L. and P.D.S.D. prepared the figures.

P.D.S.D. and J.J.L. wrote the manuscript with coauthors.

All authors reviewed and contributed to editing the manuscript.

## Acknowledgements

We thank the zebrafish research community for the numerous reagents shared and members of our laboratory for critical discussions. We thank Jesus Olvera and Cody Fine in the UCSD Human Embryonic Stem Cell Core Facility for technical assistance with flow cytometry experiments and supported by funds from the CIRM Major Facilities Grant (FA1-00607) to the Sanford Consortium for Regenerative Medicine. We thank Drs. Alexandre Colas, Gregg Duester and Alessandra Sacco for their critical reading of the manuscript and apologize to authors whose work we were not able to cite due to space constraints. This work was supported by funds from the W. M. Keck Foundation (2017-01) and National Institutes of Health (NIH) U01DK105541 to P.D.S.D., Diabetes Research Connection (DRC) Project #08 to J.J.L, and the Larry L. Hillblom Foundation Fellowship (#2016-D-008-FEL) to J.M.

## References

1 Waddington, C. H. The Strategy of the Genes. London: Allen and Unwin (1957).

2 de Lazaro, I. & Kostarelos, K. Engineering Cell Fate for Tissue Regeneration by In Vivo Transdifferentiation. Stem Cell Rev 12, 129–139, doi:10.1007/s12015-015-9624-6 (2016).

3 Riddle, M. R. et al. Transorganogenesis and transdifferentiation in C. elegans are dependent on differentiated cell identity. Dev Biol 420, 136–147, doi:10.1016/j.ydbio.2016.09.020 (2016).

4 Kalacheva, N. V., Eliseikina, M. G., Frolova, L. T. & Dolmatov, I. Y. Regeneration of the digestive system in the crinoid Himerometra robustipinna occurs by transdifferentiation of neurosecretory-like cells. PLoS One 12, e0182001, doi:10.1371/journal.pone.0182001 (2017).

5 Ariyachet, C. et al. Reprogrammed Stomach Tissue as a Renewable Source of Functional beta Cells for Blood Glucose Regulation. Cell Stem Cell 18, 410–421, doi:10.1016/j.stem.2016.01.003 (2016).

6 Halloran, M. C. et al. Laser-induced gene expression in specific cells of transgenic zebrafish. Development 127, 1953–1960 (2000).

7 Dickmeis, T. et al. A crucial component of the endoderm formation pathway, CASANOVA, is encoded by a novel sox-related gene. Genes Dev 15, 1487–1492, doi:10.1101/gad.196901 (2001).

8 Kikuchi, Y. et al. casanova encodes a novel Sox-related protein necessary and sufficient for early endoderm formation in zebrafish. Genes Dev 15, 1493–1505, doi:10.1101/gad.892301 (2001).

9 Reim, G., Mizoguchi, T., Stainier, D. Y., Kikuchi, Y. & Brand, M. The POU domain protein spg (pou2/Oct4) is essential for endoderm formation in cooperation with the HMG domain protein casanova. Dev Cell 6, 91–101 (2004).

10 Sakaguchi, T., Kikuchi, Y., Kuroiwa, A., Takeda, H. & Stainier, D. Y. The yolk syncytial layer regulates myocardial migration by influencing extracellular matrix assembly in zebrafish. Development 133, 4063–4072, doi:10.1242/dev.02581 (2006).

11 Mazzotti, A. L. & Coletti, D. The Need for a Consensus on the Locution “Central Nuclei” in Striated Muscle Myopathies. Front Physiol 7, 577, doi:10.3389/fphys.2016.00577 (2016).

12 Alexander, J. & Stainier, D. Y. A molecular pathway leading to endoderm formation in zebrafish. Curr Biol 9, 1147–1157, doi:10.1016/S0960-9822(00)80016-0 (1999).

13 Xu, Y. et al. Fast skeletal muscle-specific expression of a zebrafish myosin light chain 2 gene and characterization of its promoter by direct injection into skeletal muscle. DNA Cell Biol 18, 85–95, doi:10.1089/104454999315655 (1999).

14 Axen, K. V. & Pi-Sunyer, F. X. Long-term reversal of streptozotocin-induced diabetes in rats by intramuscular islet implantation. Transplantation 31, 439–441 (1981).

15 Capito, C. et al. Mouse muscle as an ectopic permissive site for human pancreatic development. Diabetes 62, 3479–3487, doi:10.2337/db13-0554 (2013).

16 Christoffersson, G. et al. Clinical and experimental pancreatic islet transplantation to striated muscle: establishment of a vascular system similar to that in native islets. Diabetes 59, 2569–2578, doi:10.2337/db10-0205 (2010).

17 Rafael, E. et al. Intramuscular autotransplantation of pancreatic islets in a 7-year-old child: a 2-year follow-up. American journal of transplantation: official journal of the American Society of Transplantation and the American Society of Transplant Surgeons 8, 458–462, doi:10.1111/j.1600-6143.2007.02060.x (2008).

18 Manfroid, I. et al. Reciprocal endoderm-mesoderm interactions mediated by fgf24 and fgf10 govern pancreas development. Development 134, 4011–4021 (2007).

19 Mfopou, J. K., Chen, B., Mateizel, I., Sermon, K. & Bouwens, L. Noggin, retinoids, and fibroblast growth factor regulate hepatic or pancreatic fate of human embryonic stem cells. Gastroenterology 138, 2233–2245, 2245 e2231-2214, doi:S0016-5085(10)00329-X [pii] 10.1053/j.gastro.2010.02.056 (2010).

20 Sawada, A. et al. Fgf/MAPK signalling is a crucial positional cue in somite boundary formation. Development 128, 4873–4880 (2001).

21 Buitrago-Delgado, E., Nordin, K., Rao, A., Geary, L. & LaBonne, C. NEURODEVELOPMENT. Shared regulatory programs suggest retention of blastula-stage potential in neural crest cells. Science 348, 1332–1335, doi:10.1126/science.aaa3655 (2015).

22 Christen, B., Robles, V., Raya, M., Paramonov, I. & Izpisua Belmonte, J. C. Regeneration and reprogramming compared. BMC Biol 8, 5, doi:10.1186/1741-7007-8-5 (2010).

23 Kagias, K., Ahier, A., Fischer, N. & Jarriault, S. Members of the NODE (Nanog and Oct4-associated deacetylase) complex and SOX-2 promote the initiation of a natural cellular reprogramming event in vivo. Proc Natl Acad Sci U S A 109, 6596–6601, doi:10.1073/pnas.1117031109 (2012).

24 Rossello, R. A. et al. Mammalian genes induce partially reprogrammed pluripotent stem cells in non-mammalian vertebrate and invertebrate species. Elife 2, e00036, doi:10.7554/eLife.00036 (2013).

25 Esch, D. et al. A unique Oct4 interface is crucial for reprogramming to pluripotency. Nat Cell Biol 15, 295–301, doi:10.1038/ncb2680 (2013).

26 Ladewig, J., Koch, P. & Brustle, O. Leveling Waddington: the emergence of direct programming and the loss of cell fate hierarchies. Nat Rev Mol Cell Biol 14, 225–236 (2013).

27 Jorstad, N. L. et al. Stimulation of functional neuronal regeneration from Muller glia in adult mice. Nature 548, 103–107, doi:10.1038/nature23283 (2017).

28 Cantarelli, E. & Piemonti, L. Alternative transplantation sites for pancreatic islet grafts. Current diabetes reports 11, 364–374, doi:10.1007/s11892-011-0216-9 (2011).

29 Heinrich, C., Spagnoli, F. M. & Berninger, B. In vivo reprogramming for tissue repair. Nat Cell Biol 17, 204–211, doi:10.1038/ncb3108 (2015).

30 Smith, D. K. & Zhang, C. L. Regeneration through reprogramming adult cell identity in vivo. Am J Pathol 185, 2619–2628, doi:10.1016/j.ajpath.2015.02.025 (2015).

31 Ohnishi, K. et al. Premature termination of reprogramming in vivo leads to cancer development through altered epigenetic regulation. Cell 156, 663–677, doi:10.1016/j.cell.2014.01.005 (2014).

32 Heslop, J. A. et al. Concise review: workshop review: understanding and assessing the risks of stem cell-based therapies. Stem cells translational medicine 4, 389–400, doi:10.5966/sctm.2014-0110 (2015).

33 Johannesson, B. et al. Comparable frequencies of coding mutations and loss of imprinting in human pluripotent cells derived by nuclear transfer and defined factors. Cell Stem Cell 15, 634–642, doi:10.1016/j.stem.2014.10.002 (2014).

34 Kelaini, S., Cochrane, A. & Margariti, A. Direct reprogramming of adult cells: avoiding the pluripotent state. Stem Cells Cloning 7, 19–29, doi:10.2147/SCCAA.S38006 (2014).

35 Narva, E. et al. High-resolution DNA analysis of human embryonic stem cell lines reveals culture-induced copy number changes and loss of heterozygosity. Nat Biotechnol 28, 371–377, doi:10.1038/nbt.1615 (2010).

36 Pera, M. F. Stem cells: The dark side of induced pluripotency. Nature 471, 46–47, doi:471046a [pii] 10.1038/471046a (2011).

37 Mizoguchi, T., Verkade, H., Heath, J. K., Kuroiwa, A. & Kikuchi, Y. Sdf1/Cxcr4 signaling controls the dorsal migration of endodermal cells during zebrafish gastrulation. Development 135, 2521–2529, doi:dev.020107 [pii] 10.1242/dev.020107 (2008).

38 Field, H. A., Ober, E. A., Roeser, T. & Stainier, D. Y. Formation of the digestive system in zebrafish. I. Liver morphogenesis. Dev Biol 253, 279–290 (2003).

39 Godinho, L. et al. Targeting of amacrine cell neurites to appropriate synaptic laminae in the developing zebrafish retina. Development 132, 5069–5079 (2005).

40 Park, H. C. et al. Analysis of upstream elements in the HuC promoter leads to the establishment of transgenic zebrafish with fluorescent neurons. Dev Biol 227, 279–293, doi:10.1006/dbio.2000.9898 (2000).

41 Jin, S. W., Beis, D., Mitchell, T., Chen, J. N. & Stainier, D. Y. Cellular and molecular analyses of vascular tube and lumen formation in zebrafish. Development 132, 5199–5209, doi:10.1242/dev.02087 (2005).

42 Roman, B. L. et al. Disruption of acvrl1 increases endothelial cell number in zebrafish cranial vessels. Development 129, 3009–3019 (2002).

43 Lancman, J. J. et al. Specification of hepatopancreas progenitors in zebrafish by hnf1ba and wnt2bb. Development 140, 2669–2679, doi:10.1242/dev.090993 (2013).

44 Clements, W. K. & Kimelman, D. LZIC regulates neuronal survival during zebrafish development. Dev Biol 283, 322–334, doi:10.1016/j.ydbio.2005.04.026 (2005).

45 Brend, T. & Holley, S. A. Zebrafish whole mount high-resolution double fluorescent in situ hybridization. J Vis Exp, doi:10.3791/1229 (2009).

46 Schoenebeck, J. J., Keegan, B. R. & Yelon, D. Vessel and blood specification override cardiac potential in anterior mesoderm. Dev Cell 13, 254–267, doi:10.1016/j.devcel.2007.05.012 (2007).

47 Zhou, Y. et al. Latent TGF-beta binding protein 3 identifies a second heart field in zebrafish. Nature 474, 645–648, doi:10.1038/nature10094 (2011).

48 Yelon, D. et al. The bHLH transcription factor hand2 plays parallel roles in zebrafish heart and pectoral fin development. Development 127, 2573–2582 (2000).

49 Zeng, X. X. & Yelon, D. Cadm4 restricts the production of cardiac outflow tract progenitor cells. Cell Rep 7, 951–960, doi:10.1016/j.celrep.2014.04.013 (2014).

50 Carey, B. W. et al. Reprogramming of murine and human somatic cells using a single polycistronic vector. Proc Natl Acad Sci U S A 106, 157–162, doi:10.1073/pnas.0811426106 (2009).

