## Supplementary material for "*In vivo* lineage conversion of vertebrate muscle into early endoderm-like cells": Combined Supplement Figures and Methods

Supplementary Figure 1 Co-expression of Oct4 and Sox32 specifically induces expression of the early endoderm sox17-GFP in non-endoderm cells.

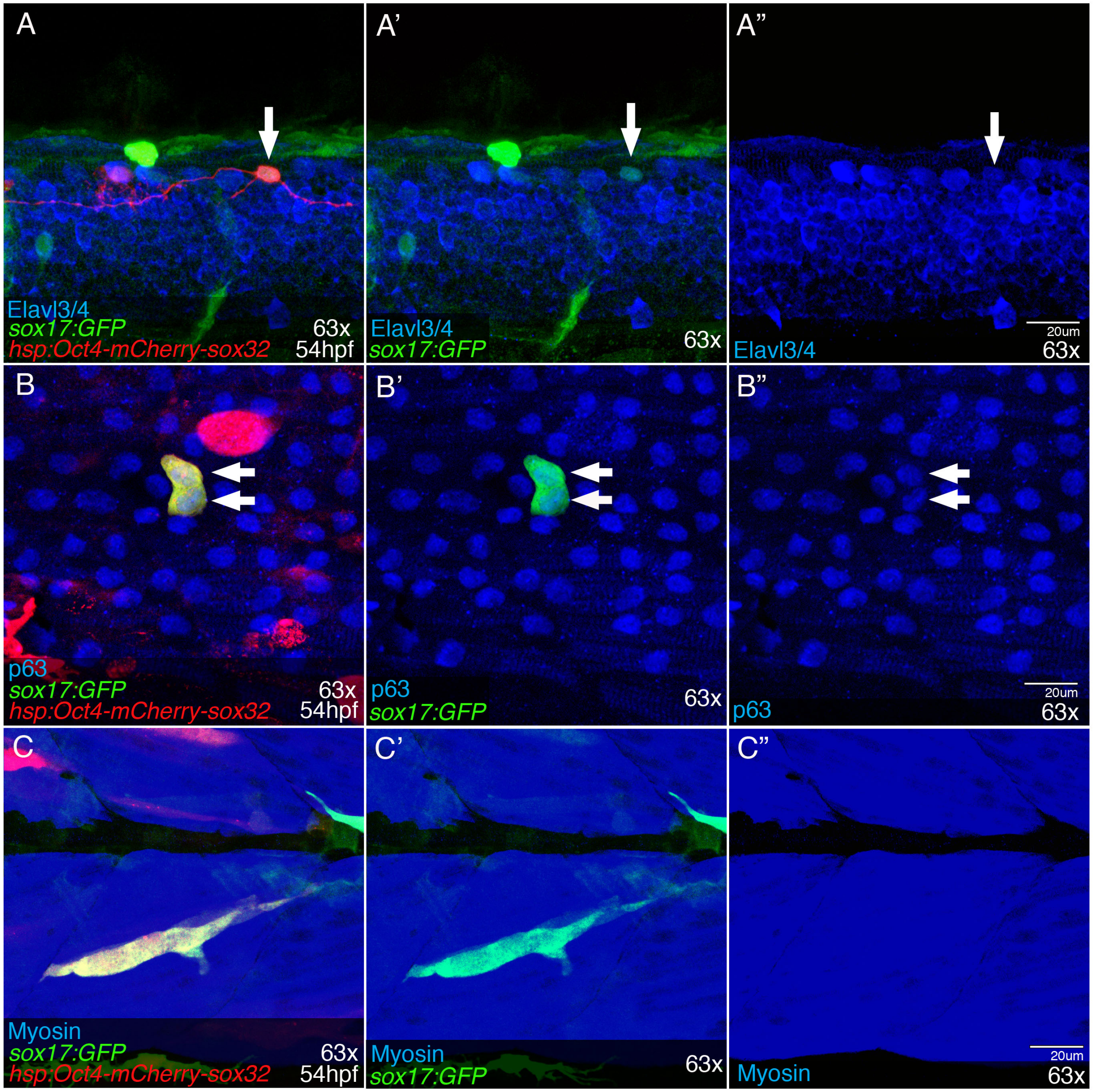

**Supplemental Figure 1: Co-expression of Oct4 and Sox32 specifically induces expression of the early endoderm sox17-GFP in non-endoderm cells.**

**(A-C”)**. Co-expression of Oct4 and Sox32 is sufficient to induce sox17:GFP in non-endoderm cells arising from ectoderm or mesoderm lineages where sox17:GFP expression is not normally detected. At 48hpf, Z-focal plane showing that coexpression of Oct4 and sox32 (red) can induce sox17:GFP (green) in neural cells marked by the early, pan-neuronal marker elavl3/4 **(blue; arrow, A-A”)** and in keratinocytes marked by p63 **(blue; arrows, B-B”)** as well as in myosin stained myocytes **(blue; C-C”)**. Note that neural cells and keratinocytes arise from the ectoderm, and myocytes from the mesoderm lineage.

Supplementary Figure 2 Co-expression of Oct4 and Sox32 does not induce vascular or neural genetic programs.

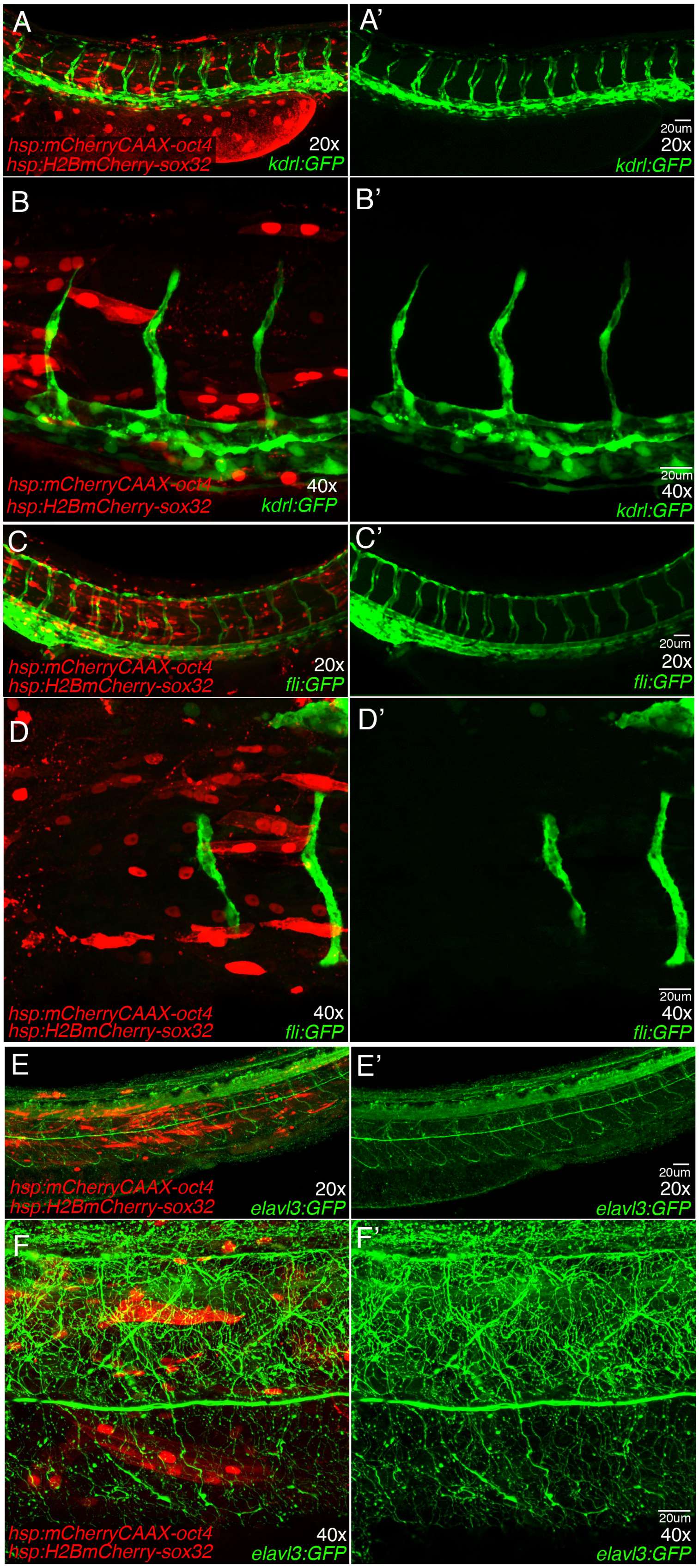

**Supplemental Figure 2: Co-expression of Oct4 and Sox32 does not induce vascular or neural genetic programs.**

**(A-F')** Z-focal plane of 48hpf zebrafish showing that cells co-expressing Oct4 (red membrane)and sox32 (red nuclei) do not up regulate detectable levels of vascular markers *kdrl:GFP* **(green A-B')** or *fli1:GFP* **(green C-D')** outside of their endogenous expression domains. **(E-F')** Moreover, co-expression of Oct4 and sox32 do not upregulate the early neural differentiation marker *elavl3:GFP.* Together, these data suggest that, following co-expression of Oct4 and sox32, muscle cells primarily up-regulate factors required for endoderm development.

Supplement Figure 3 Nuclei aggregation in reprogrammed muscle cells

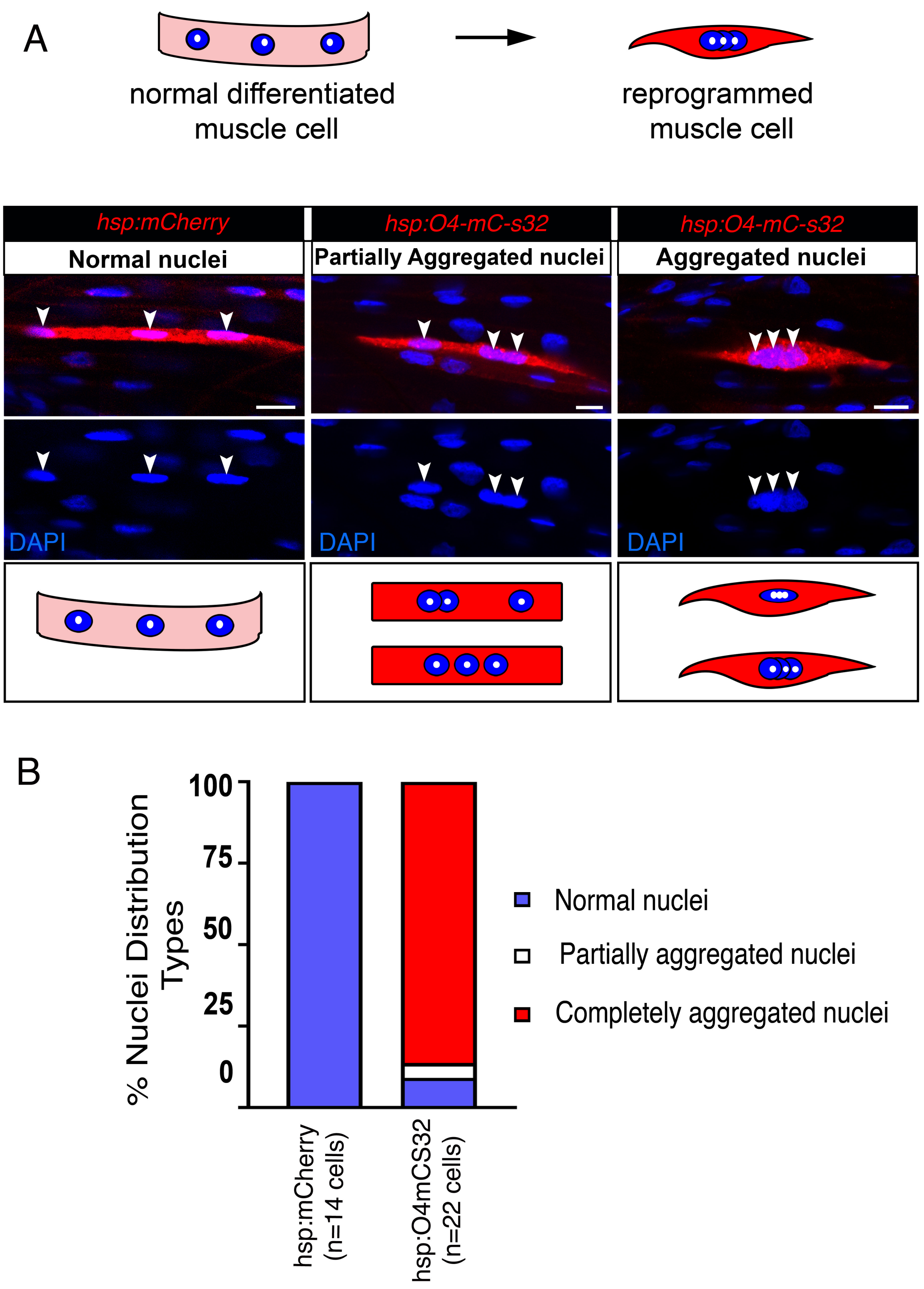

**Supplement Figure 3:** Nuclei aggregation in reprogrammed muscle cells. **(A, Top**) Diagram depicting nuclei **(blue, nuceolus-white)** position in normal 54 hpf zebrafish muscle cells **(pink)** and in reprogrammed muscle cells **(red). (A, Bottom**) Representative confocal Z-stack images of 54hpf zebrafish muscle cells expressing mCherry alone (red; left-control) or Oct4-mCherry-sox32 **(red; middle, right**) with DAPI stained nuclei **(blue).** The nuclei in control muscle cells **(left; *hsp:mCherry*)** are regularly positioned along the length of the muscle cell whereas in reprogrammed muscle cells, the nuceli aggregate either partially **(middle)** or completely **(right)** in the center of the cell. **(B)** Bar graph displaying incidence of nuclear aggregation in *hsp:mCherry* positive muscle cells **(control; 0/19 cells with nucelar aggregation) and** reprogrammed *hsp:Oct4-mCherry-sox32* mucle cells **(2/22 no aggregation, 1/22 partial aggregation, 19/22 completely aggregated).**

Supplement Figure 4   Loss of myh7 and Myosin in reprogramed muscle cells.

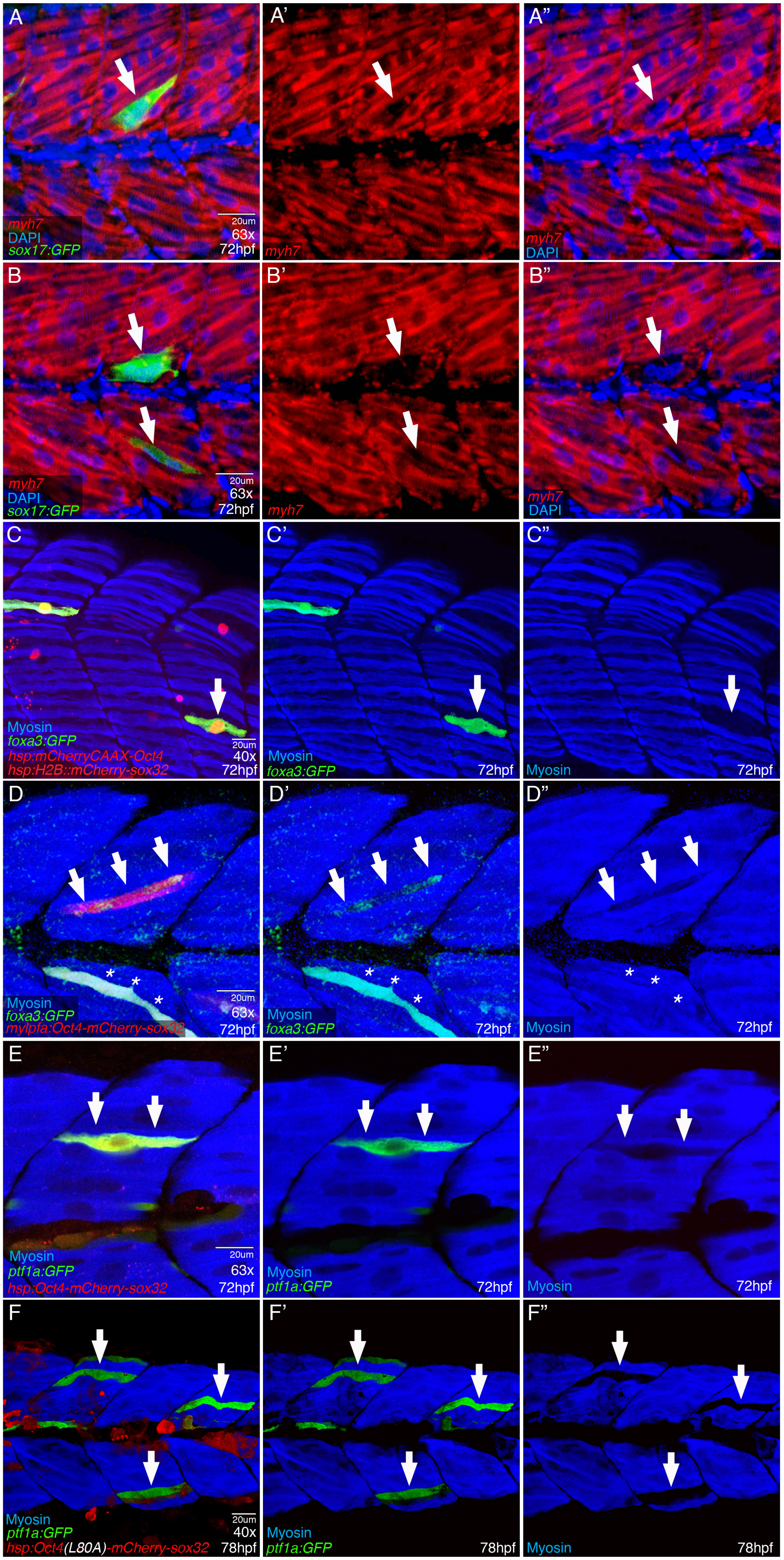

**Supplement Figure 4**: **Loss of *myh7* and Myosin in reprogramed muscle cells.**
**(A-B”)** 3D rendering of representative 48hpf zebrafish with flourescent *whole mount in situ hybridization* and immunohistochemistry to detect *myosin heavy chain 7* mRNA **(*myh7*; red)** and sox17:GFP **(green)** in myocytes from *hsp:Oct4-P2A-mCherry-P2A-sox32* injected embryos. Arrows highlight muscle cells that have up regulated sox17:GFP **(green)** and subsequently down-regulated *myh7* **(red; n=11/13 reprogrammed cells (3 independent samples) )**. Also note that nuclei in reprogrammed myocytes appear to aggregate and are no longer regularly positioned along the horizontal axis as they are in neighboring, unaffected myocytes. **(C-F”)** 3D rendering of 72hpf zebrafish with Myosin stained myocytes **(blue)** injected with various Oct4 and sox32 expression constructs. **(C-C”)** Example of a single myocte coexpressing mCherryCAAX-P2A-Oct4 and H2B::mCherry-P2A-sox32 with induced foxa3:GFP **(green) (C, C’; arrow)** that has also lost expression of myosin **(blue; C”; arow)**. **(D-D”)** The muscle specific construct *mylpfa:Oct4-P2A-mCherry-P2A-sox32* was used to reprogram muscle cells **(D, arrows, red)**, resulting in upregualtion of foxa3:GFP **(D’, arrow; green)** and loss of myosin **(D”, arrows; blue)**. Not all reprogrammed muscle cells that up regulate foxa3:GFP in the same animal lose myosin (D-D”; asteriks). (Note: This image is the same as that used in Figure 2F, F’.) **(E-E”)** *hsp:Oct4-P2A-mCherry-P2A-sox32* was used to reprogram muslce cells in *ptf1a:GFP* transgenic zebrafish. *ptf1a:GFP* **(green; E, E’, double arrows)** is induced in a myocte that has also lost expression of myosin **(E”, blue; double arrow). (F-F”)** Myoctes reprogrammed with the pluripotent defective Oct4(L80A) mutant and sox32 up regulate ptf1a:GFP **(green; F, F’, arrows)** and can also lose Myosin **(blue; F”, arrows).** (Note: This image is the same image used in Figure 4D-D”).

Supplementay Figure 5 A time course of endoderm gene induction in reprogrammed muscle cells.

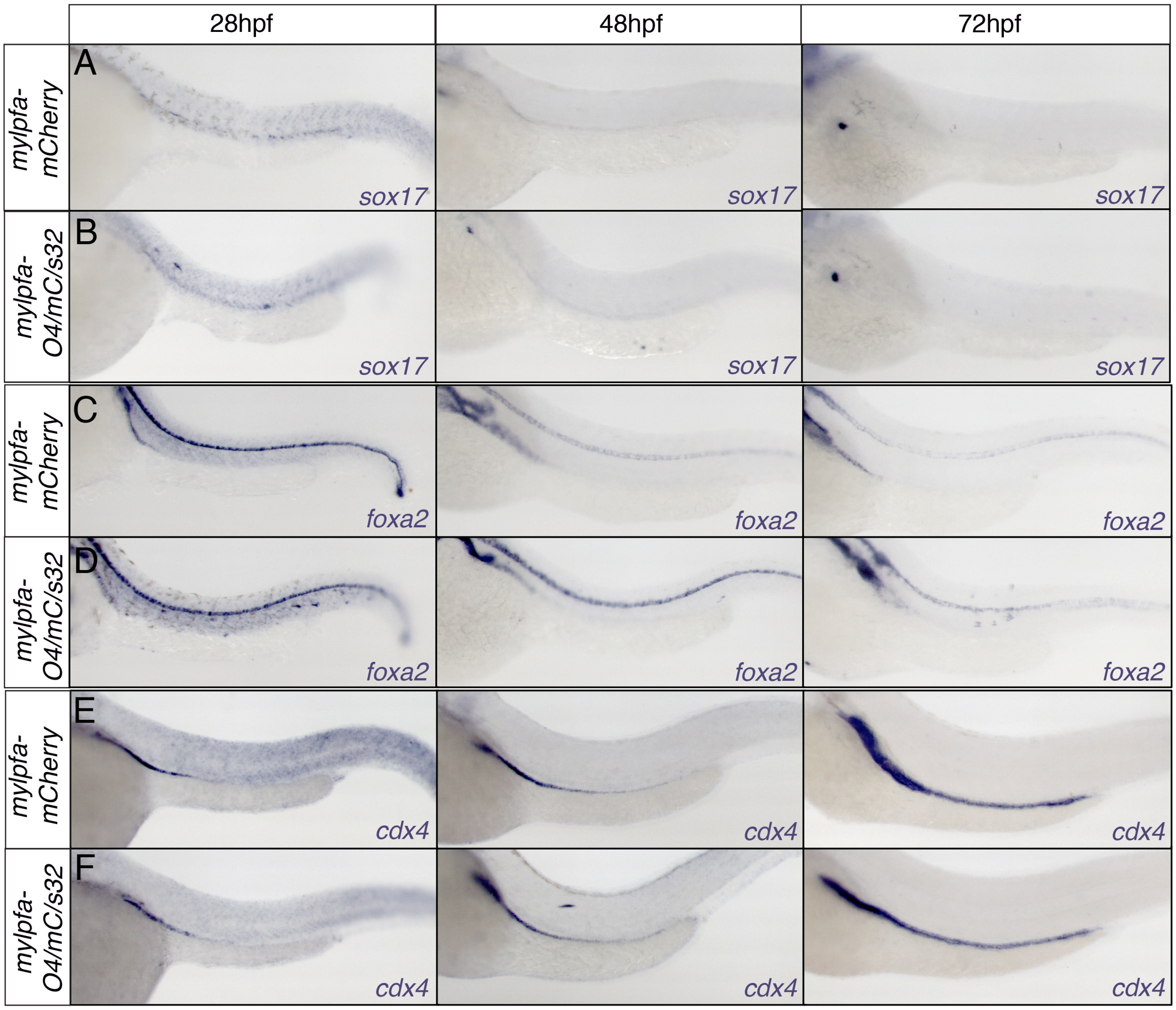

**Supplement Figure 5:** A time course of endoderm gene induction in reprogrammed muscle cells. **(A-F)** Whole mount *in situ hybridization* of 28hpf **(left column)**, 48hpf **(middle column)** and 72hpf zebrafish embryos **(right column)** injected with either the muscle specific control construct *mylpfa:mCherry* **(control; A, C, E)** or *mylpfa:Oct4-P2A-mCherry-P2A-sox32* **(B, D, F)**. Compared to mylpfa:mCherry controls **(A)**, ectopic *sox17* expression (purple)can be detected within the myotomes at 28hpf, but is not detected at 48hpf or 72hpf **(arrows; B)**. **(C-D)** Embryos were probed for ectopic *foxa2* expression (purple) which is not detected outside its endogenous domain in control samples **(C)** but is detected in the trunk region at 28hpf, 48hpf and 72hpf following coexpression of Oct4 and sox32. **(E-F)** In mylpfa:mCherry injected controls, *cdx 4* expression (purple) is never detected outside its endogenous domain **(E)** but in experimental embryos, ectopic *cdx4* expression can be observed within the myotomes at 48hpf but it is not detected at 28hpf or 72hpf.

Supplement Figure 6 Reprogrammed muscle cells produce extensions and display drastic changes to cell body morphology.

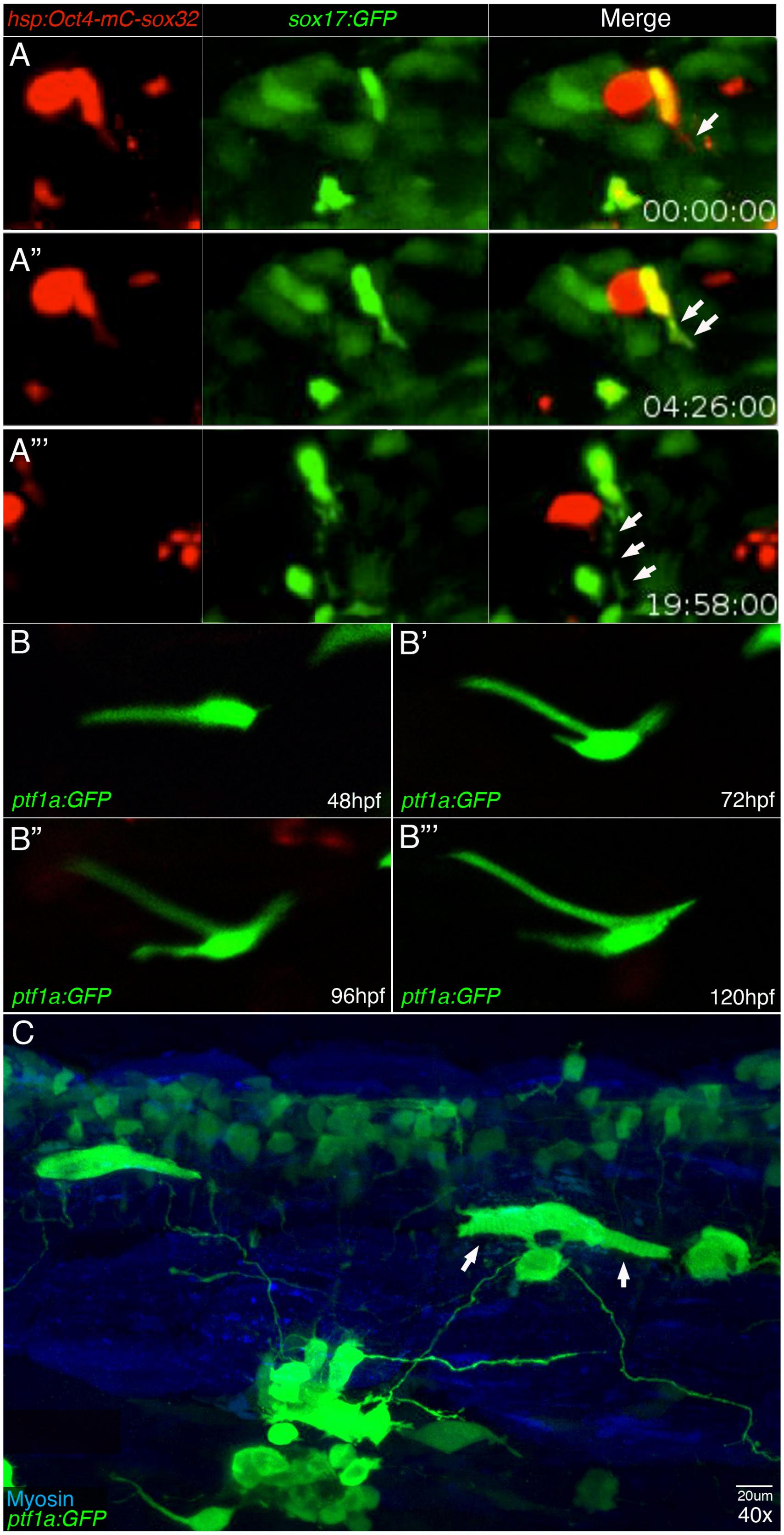

**Supplement Figure 6: Reprogrammed muscle cells produce extensions and display drastic changes to cell body morphology.**

**(A-A”)** Stills captured from live imaging movies of sox17:GFP transgenic zebrafish injected with *hsp:Oct4-P2A-mCherry-P2A-sox32* and imaged from 48-72hpf using light sheet confocal microscopy. **(A)** Reprogrammed cell **(yellow arrowhead)** residing in the head region, up regulates sox17:GFP **(green)** and produces a long cell extension over 20 hours **(A; single arrow, A”; double arrow, A”’; triple arrow). (B-B”’)** Live confocal imaging time course of a reprogrammed muscle cell that unregulated ptf1a:GFP **(green; B-B’”’)** and tracked over 5 days. During this time, cell morphology is radically altered as long extensions are extended from multiple locations along the main cell body. **(C)** 3D rendering of a 72hpf ptf1a:GFP transgenic zebrafish trunk injected with *hp:Oct4-P2A-mCherry-P2A-sox32*. Multiple cells with ectopic ptf1a:GFP **(green, arrows)** produce long projections and potentially contact other distant reprogrammed ptf1a:GFP expressing cells.

### Antibody Details

Table 1

| Primary Antibody | | Company | Part# | Concentration | | |
|---|---|---|---|---|---|---|
| **GFP** | Anti-Chicken | Aves Labs | GFP-1020 | 1:300 | Chicken | |
| **Insulin** | Anti-Guinea Pig | biomeda | V2024 | 1:200 | Guinea Pig | |
| **mCherry** | Anti-Rabbit | Rockland | 600-401-P16S | 1:200 | Rabbit | |
| **Myosin Heavy Chain (F-59)*** | Anti-Mouse | Developmental Studies Hybridoma Bank | F-59 | 1:20 | Mouse | |
| **myosin light chain 1 and 3f (LC1f/3f: F310)*** | Anti-Mouse | Developmental Studies Hybridoma Bank | F310 | 1:20 | Mouse | |
| **HuC/D (Elavl3/4)** | Anti-Rabbit | abcam | ab210554 | 1:100-1:500 | Rabbit | |
| **Tp63** | Anti-Rabbit | GeneTex | GTX124660 | 1:200 | Rabbit | |
| ***The monoclonal antibodies F59 and F310 developed by F.E. Stickdale was obtained from the Developmental Studies Hybridoma Bank, created by the NICHD of the NIH and maintained at The University of Iowa, Department of Biology, Iowa City, IA 52242.** | | | | | | |
| **Secondary Antibody/Stains** | | | | | | |
| **AlexaFluor 488** | Anti-Chicken | Jackson ImmunoResearch | 703-545-155 | | | Donkey |
| **DyLight 405** | Anti-Guinea Pig | Jackson ImmunoResearch | 706-475-148 | 1:200 | | Donkey |
| **AlexaFlour 647** | Anti-Mouse | Invitrogen | | 1:200 | | Donkey |
| **AlexaFluor 594** | Anti-Rabbit | Invitrogen | | 1:200 | | Goat |
| **AlexaFluor 594** | Anti-Rabbit | Invitrogen | | 1:200 | | Goat |
| **AlexaFluor 568** | Anti-Rabbit | Invitrogen | | 1:200 | | Donkey |
| **AlexaFluor 594** | Anti-Chick | Invitrogen | | 1:200 | | Donkey |
| **DAPI (500mg/ml)** | | Invitrogen | | 1:200 | | |

### qPCR Primers used in study

| Primer | Forward | Reverse |
|---|---|---|
| ef1a | AGAAGGCTGCCAAGACCAAG | AGAGGTTGGGAAGAACACGC |
| sox17 | GCATCCGAAGGCCAATGAAC | GCTTTCCATGACTTACCAAGC |
| foxA2 | TCGTGTGGGGAAGCGTTTTA | CGAGGTGTAACACTCAGGCT |
| foxA3 | GGGATGTTGAGCTCCGTGAA | CGGAGAGGAATACATCTCATTTGC |
| pdx1 | ACCATCTCCCATTTCCGTGG | TCGACCATATAAGGGCCTGTC |
| ptf1a | ACCGAGGAACAAGATCCCCAT | CCAGACTTTCGCTGTCCGAA |
| cdx1a | CACGGACGAAGGACAAGTACA | GATCTTGACCTGGCGTTCTGA |
| hnf1ba | ATGTTCCCACTGCCATTGCT | ACAATGTGGAACAAATCACATCTTG |
| nkx6.1 | CTCGCTATCCCAAACCCCTG | TTTTGATGTGGTGAGCACGC |
| nkx2.2 | CGCCTGGAGTGTTAGTGCAA | GGACAGGCCGTGTAATGAGT |
| hnf4a | GAGCACAGACTCCTCACCAC | TAGAGTGCCTGCCCCTAAGT |
| tnnt3b | GAGGTAGAGGTAGCCCCAGA | ACTGCAACTACTGCTAAGACCA |
| mylpfa | GAGGGTTCCTCCAACGTCTT | GAGCTCGGCATCGCTTTTAG |
| myoD | CACACCAAATGCTGACGCAC | TGTGGAAATTCGCTCCACGA |
| acta1 | ACGATGATGAGACCACAGCTT | TACCAACCATCACACCCTGG |
| myhz2 | CAAGGAACGCAAGTAAGCCG | ACAAGCGGTTTTGGCATCAA |
| tnnc1a | TGAAGATGGAAGTGGTACGGTG | CATACGGAAGAGTTCCGCCA |
| pax3a | CAGCAAACCCAAGCAGAGCAC | TCCGATCGCAGATTCCATCTTT |
| pax7a | TGAATCCTGTGAGCAACGGC | TGCTCTTGATCTGTGAAGCGT |
| meox1 | ACCTCACTGAGAGACAGGTGA | TGCTTCAAGGTCGTGAGGAG |
| fli1a | GTCTCTCCGCCACATATCGG | ACTGACAGCGCCTCCTTAAT |
| kdrl | TCCCATTGAAAACGTTGATGACC | TAGCTGTTTTCACCACCAGGG |
| tbxta | CCAACACCAGTCAGTACCCA | CATCGAAGAACCGCGTAGGA |
| zic2.2 | CACATGAAGGTTCACGAGGA | CCGAGCATGGAGAGATCAGAC |
| sox19b | CGCCAGCTCTTACAGTCAAATG | ACGGTGGTGGTTTGGTACTC |
| foxi1 | GGATGATCCTGGGAAAGGAAAT | CCAATTTAAGCGCGTCCTCG |
| p63 | GCTCGGCCTGTTTGGACTAT | TCAGCCTGGACAAGTCCTCTA |
| oct4 (endogenous) | CCAATGGGAGAGAAGTTGGT | GATTGCGCGTCTCAGTATCA |
| myca | TATGCTGCAAGTGACCGGAG | TCACCGGCATTTTGACACTTG |
| nanog | AAGACTGAGCCCGACCAAAA | AGCTCCAGGAATCTGGCGT |

| | | |
|---|---|---|
| vasa | GGAGGAAGATCAGAGTCCCG | TCCATCAGAACCATTTGAGCCT |

| sox17 | mylpfa-mCherry | mylpfa-mO/mC/s32 |
|---|---|---|
| | 6 | 34.25 |
| | 5.9 | 26.25 |

| elavl3 | mylpfa-mCherry | mylpfa-mO/mC/s32 |
|---|---|---|
| | 0.6 | 1.3 |
| | 0.38 | 0 |

| fli1a | mylpfa-mCherry | mylpfa-mO/mC/s32 |
|---|---|---|
| | 7 | 3.05 |
| | 8.3 | 7.05 |

Efficieny of induction statistics and graph

| | |
|---|---|
| Table Analyzed | sox17 |
| Column A | mylpfa-mCherry |
| vs. | vs. |
| Column B | mylpfa-mO/mC/s32 |
| Unpaired t test | |
| P value | 0.026 |
| P value summary | * |
| Significantly different (P < 0.05) | Yes |
| One- or two-tailed P value? | Two-tailed |
| t, df | t=6.075, df=2 |
| How big is the difference? | |
| Mean of column A | 5.95 |
| Mean of column B | 30.25 |
| Difference between means (A - | -24.30 ± 4.000 |
| 95% confidence interval | -41.51 to -7.088 |
| R squared (eta squared) | 0.9486 |
| F test to compare variances | |
| F, DFn, Dfd | |
| P value | |
| P value summary | |
| Significantly different (P < 0.05)? | |
| Data analyzed | |
| Sample size, column A | 2 |
| Sample size, column B | 2 |

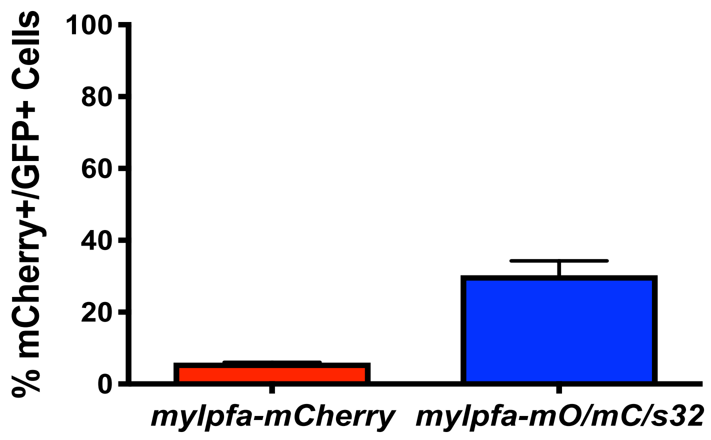

Table Analy huC

| | |
|---|---|
| Column A | mylpfa-mCherry |
| vs. | vs. |
| Column B | mylpfa-mO/mC/s32 |

| | |
|---|---|
| Unpaired t test | |
| P value | 0.8309 |
| P value sun | ns |
| Significantl | No |
| One- or two | Two-tailed |
| t, df | t=0.2427, df=2 |

| | |
|---|---|
| How big is the difference? | |
| Mean of col | 0.49 |
| Mean of col | 0.65 |
| Difference k | -0.1600 ± 0.6592 |
| 95% confid | -2.996 to 2.676 |
| R squared ( | 0.02861 |

F test to compare variances
F, DFn, Dfd
P value
P value summary
Significantly different (P < 0.05)?

| | |
|---|---|
| Data analyzed | |
| Sample size | 2 |
| Sample size | 2 |

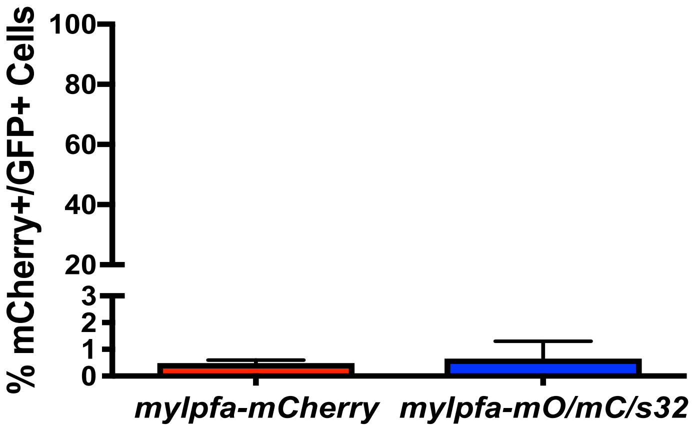

| | |
|---|---|
| Table Analy | fli1a |
| Column A | mylpfa-mCherry |
| vs. | vs. |
| Column B | mylpfa-mO/mC/s32 |
| Unpaired t test | |
| P value | 0.3418 |
| P value sun | ns |
| Significantl | No |
| One- or two | Two-tailed |
| t, df | t=1.236, df=2 |
| How big is the difference? | |
| Mean of col | 7.65 |
| Mean of col | 5.05 |
| Difference k | 2.600 ± 2.103 |
| 95% confid | -6.448 to 11.65 |
| R squared ( | 0.4332 |
| F test to compare variances | |
| F, DFn, Dfd | |
| P value | |
| P value summary | |
| Significantly different (P < 0.05)? | |
| Data analyzed | |
| Sample size | 2 |
| Sample size | 2 |

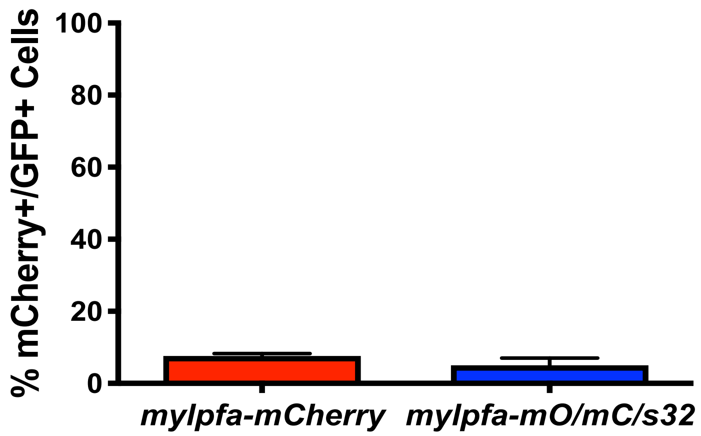

Flow Cytometry Gating Strategy

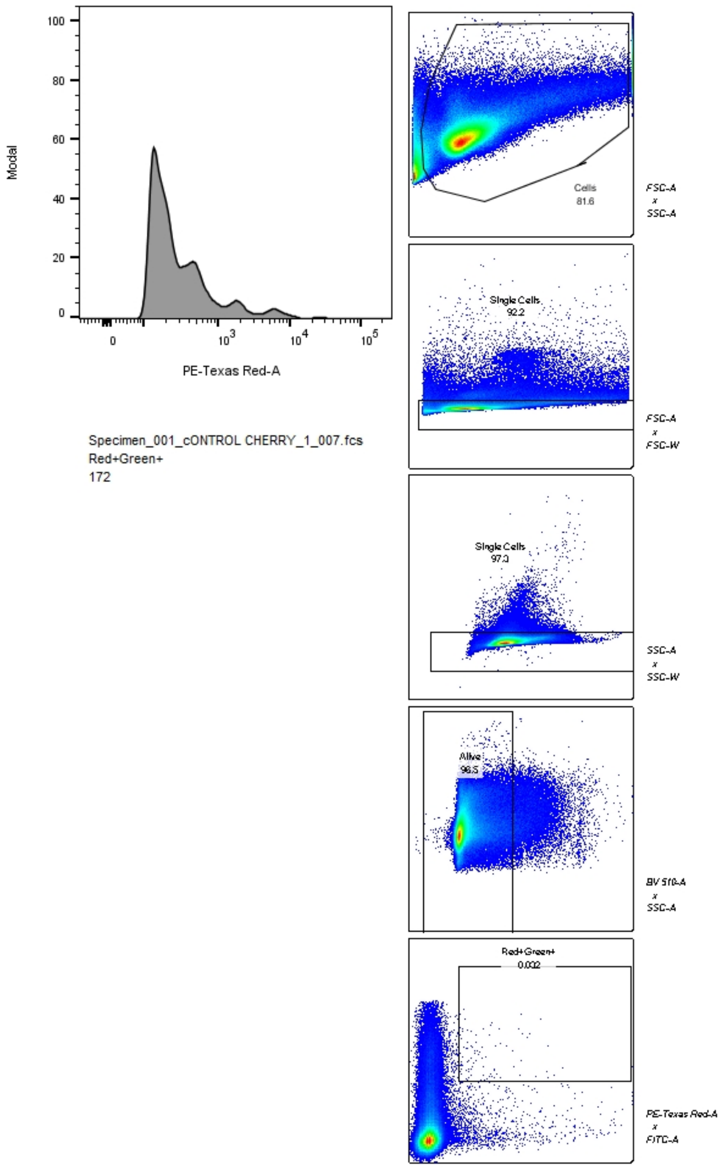

Flow cytometry: FSC-A x SSC-A to gate on all cells; FSC-A x FSC-W to gate on single cells; SSC-A x SSC-W to gate on single cells; BV510-A x SSC-A to gate on alive cells; PE-TexasRed-A x FITC-A to gate on green+ and red+ cells

### FACS Gating Strategy

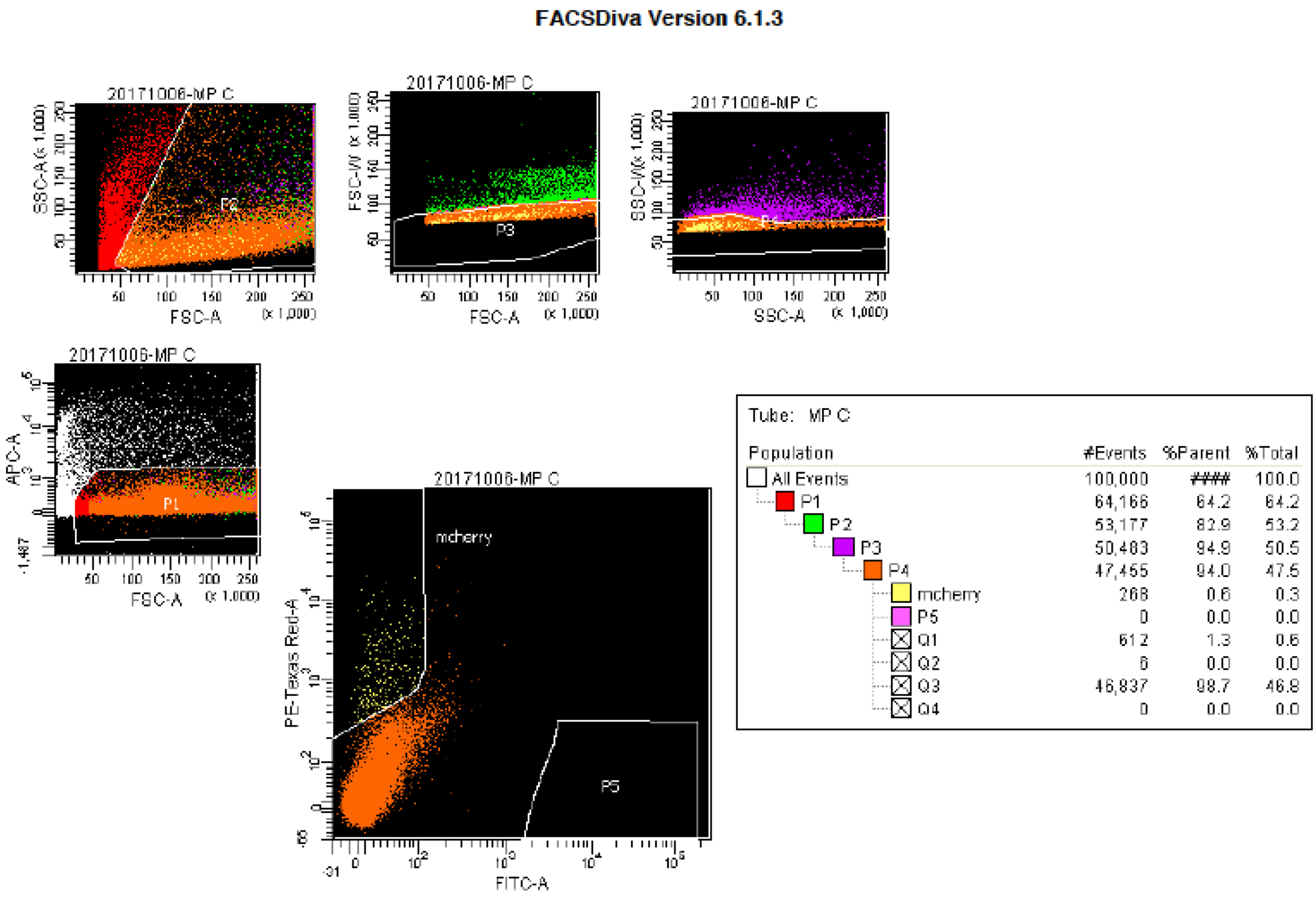

Tube: MP C

| Population | #Events | %Parent | %Total |
|---|---|---|---|
| All Events | 100,000 | #### | 100.0 |
| P1 | 64,166 | 64.2 | 64.2 |
| P2 | 53,177 | 82.9 | 53.2 |
| P3 | 50,483 | 94.9 | 50.5 |
| P4 | 47,465 | 94.0 | 47.5 |
| mcherry | 268 | 0.6 | 0.3 |
| P5 | 0 | 0.0 | 0.0 |
| Q1 | 612 | 1.3 | 0.6 |
| Q2 | 6 | 0.0 | 0.0 |
| Q3 | 46,837 | 98.7 | 46.8 |
| Q4 | 0 | 0.0 | 0.0 |

FACS: FSC-A x SSC-A to gate on all cells; FSC-A x FSC-W to gate on single cells; SSC-A x SSC-W to gate on single cells; APC-A x FSC-A to gate on alive cells; PE-TexasRed-A x FITC-A to gate red+ cells
